# Regulated mRNA recruitment in dinoflagellates is reflected in hyper-variable mRNA spliced leaders and novel eIF4Es

**DOI:** 10.1101/2024.02.20.581179

**Authors:** Grant D. Jones, Ernest P. Williams, Saddef Haq, Tsvetan R. Bachvaroff, M. Basanta Sanchez, Allen R. Place, Rosemary Jagus

## Abstract

Dinoflagellates are eukaryotic algae with large genomes that rely heavily on post-transcriptional control for the regulation of gene expression. Dinoflagellate mRNAs are *trans*-spliced with a conserved 22 base spliced leader sequence (SL) that includes the 5’-cap to which the translation initiation factor 4E (eIF4E) binds to facilitate ribosomal recruitment. The binding of an eIF4E to a specific mRNA SL is a potential regulatory point in controlling dinoflagellate gene expression. Here we show that m^7^G is the 5’-cap base of the 65 bp SL RNA with additional methylations throughout the SL to give a mixture of novel multi-methylated sequences in *Amphidinium carterae* (CCMP1314). There is also sequence variability in all four bases seen at the first position followed by a variety of polymorphisms. Three novel clades of eIF4E have been shown in dinoflagellates that are distinct from the three metazoan classes of eIF4E. Members of each clade differ significantly from each other, but all bear the distinctive features of a cap-binding protein. Here we show large differences in expression and activity in six of the eight eIF4E family members from *A. carterae*. Transcripts of each are expressed throughout the diel cycle, but only eIF4E-1 family members and eIF4E-2a show discernable expression at the level of protein. Recombinant eIF4E-1 family members and eIF4E-3a, but not eIF4E-2a, are able to bind to m^7^GTP substrates *in vitro*. Overall, eIF4E-1a emerges with characteristics consistent with the role of a prototypical initiation factor; eIF4E-1a is the most conserved and highly expressed eIF4E family member, has the highest affinity for m^7^GpppG and m^7^GpppC by surface plasmon resonance, and is able to complement a yeast strain conditionally deficient in eIF4E. The large number of eIF4E family members along with the sequence and methylation state variability in the mRNA SLs underscore the unique nature of the translational machinery in the dinoflagellate lineage and suggest a wide range of possibilities for differential recruitment of mRNAs to the translation machinery.

**Impact Statement:** *In the dinoflagellate, <u>A. carterae</u>, hyper-variable mRNA spliced leaders and novel eIF4Es reflect the reliance of dinoflagellates on variable mRNA recruitment for the regulation of gene expression*.

## Introduction

Dinoflagellates are ecologically important microbial eukaryotes of marine and freshwater habitats that play a crucial role in food webs as both primary producers and consumers (1–5). They are known for the formation of toxic blooms in coastal waters (6) for providing the most abundant source of bioluminescence in the ocean (7), and for being essential symbionts of corals (8). Dinoflagellates are members of the superphylum Alveolata, which also includes ciliates and apicomplexans. Evidence from phylotranscriptomic analyses suggest that core dinoflagellates emerged at the Permian/Triassic boundary (9). Dinoflagellate relationships are now well resolved; initially, the deeper branches of the lineage came into focus with the use of ribosomal proteins (10). With expanded transcriptome datasets, using well defined conserved eukaryotic proteins, a strongly supported dinoflagellate phylogeny has emerged (9).

Dinoflagellate genomes have many divergent genetic traits from well-studied model eukaryotes. Their genomes are extremely large, with up to 200 pg DNA per nucleus (11, 12). In addition, duplicate gene copies, often large in number, are arranged in tandem arrays (13, 14) that may, or may not, be polycistronically transcribed (15, 16) and there is a reduced role for transcriptional regulation compared to other eukaryotes (17–20). This leaves many questions concerning how dinoflagellates regulate their gene expression and respond to environmental stresses.

Overall, the degree to which dinoflagellates use transcriptional responses to alter gene expression appears limited (15, 19–21) and there is only a slight correspondence between mRNA expression levels and protein expression (18, 22, 23). However, the proteome has been shown to change readily in response to classic stressors, as well as to diurnal changes (22, 24–26). A benchmark for dinoflagellate proteome changes is seen in the bioluminescence of *Lingulodinium polyedrum* (27). The mRNA level of the luciferase binding protein is maintained throughout a diel cycle, while protein abundance increases only at night and decreases to non-detectable levels during the day. Overall, many such studies have pointed to the regulation of mRNA recruitment as being important in dinoflagellate gene expression.

mRNAs in dinoflagellates are *trans*-spliced with an apparently ubiquitous 22 nucleotide sequence termed the spliced leader (SL) derived from a spliced leader RNA (SL RNA) (15, 28). In spliced leader *trans*-splicing, the exonic portion of the spliced leader transcript is transferred to the 5’-end of coding transcripts. to yield a mature mRNA (reviewed (29)). This process adds the SL sequence, including the RNA cap structure, to the 5’ end of coding transcripts. Spliced leader *trans*-splicing is found in many unrelated eukaryotic lineages; in protists such as euglenozoans, perkinsozoans, dinoflagellates, and in many animal branches such as cnidarians, platyhelminths, nematodes and some chordates (30, 31). Although both the mRNA cap structure and the SL are likely to impact mRNA recruitment processes, investigations into these have not been rigorously pursued in dinoflagellates. Our earlier findings of a large family of sequences encoding the translational initiation factor eIF4E (mRNA cap-binding protein) in dinoflagellates also imply the importance of mRNA recruitment in the regulation of gene expression in dinoflagellates (32).

Dinoflagellate genomes are represented by draft short-read assemblies of the coral reef symbionts, *Symbiodinium B-1* and *Symbiodinium kawagutii* (33, 34) and others (35) and two strains of the closely related, but free-living psychrophile, *Polarella glacialis* (14). These are great resources for the dinoflagellate community, but they represent the small genomes of purely symbiotic and/or highly specialized lineages and therefore may not encompass the full set of gene diversity and life strategies present in the overall diversity of dinoflagellates. However, a wealth of sequence data is available from the transcriptomic datasets collected under the Marine Microbial Eukaryote Transcriptome Sequencing Project (MMETSP), http://marinemicroeukaryotes.org/). The transcriptome data provides a resource for performing comprehensive analysis of multiple dinoflagellate species which we have used to identify the core translational machinery of dinoflagellates (32, 36, 37), with the anticipation that this will lead to a better understanding of their biology with respect to gene expression.

The present study focuses on key features of dinoflagellate mRNA recruitment as a point of control and in particular the 5’-cap base, the spliced leader (SL) and the family of eIF4E translation factors. Translation initiation is a coordinated rate-limiting process involving many components that function to recruit mRNA to the protein translation machinery (reviewed in (38)). This multistep process has been relatively well-defined in model systems such as the yeast, *S. cerevisiae*, and mouse/human. mRNA recruitment involves the binding of the translation initiation factor eIF4E to the 7-methylguanosine cap structure (m7G) at the 5’-end of mRNA (reviewed in (38)). eIF4E also interacts with a scaffold protein eIF4G which, in turn, interacts with additional initiation factors that enable recruitment of mRNA by the ribosomal 43S preinitiation complex.

Through phylogenetic analyses, eIF4E has been shown to be part of an extended gene family of orthologues and paralogues found exclusively in eukaryotes (Hernández et al., 2016, #51420, 36, 39–42). Expansion of the eIF4E family has occurred across eukaryotes, from excavates (such as the trypanosomatids) to the different multicellular lineages. Multiple eIF4E family members have been identified in a wide range of organisms that include plants, fungi, metazoans, and various protist lineages (36, 38–41, 43–46). In metazoans, some eIF4E family members can support global recruitment of mRNAs, others promote or inhibit translation of specific mRNAs or subsets of mRNAs, or play a role in remediation of stalled ribosomes. Some eIF4E family members are involved in regulation of translation in response to stress, others show variable affinities for different cap structures, different eIF4Gs, or may not be involved in translation at all but in a range of processes, including mRNA processing, export, translation, storage, and decay (38).

In plants, fungi and metazoans, eIF4E family members form three classes; Class I, Class II and Class III, with Class I containing the canonical cap binding translation factors. This classification was based on variations in the residues equivalent to Trp-43 and Trp-56 of human eF4E (40). Our earlier survey of the eIF4E family in protists showed a wide diversity and number of orthologs/paralogs (36). eIF4E sequences from dinoflagellates, ciliates, heterokonts (32, 36, 37), and the kinetoplastids *Trypanosoma* and *Leishmania* (47) do not correspond to this classification. Different strategies have evolved during evolution to accommodate cap-dependent translation for differing requirements, involving various forms of eIF4E not encountered across the eukaryotic domain overall (41). For instance, in protozoan parasites of the Trypanosomatidae family, a unique cap4-structure is located at the 5’-UTR of all mRNAs. Six different eIF4E family members and five eIF4Gs are active depending on the life cycle stage of the parasite (47). Even more striking is the number and diversity of eIF4Es found in the dinoflagellate. Analysis of the transcriptomes of eleven dinoflagellate species has established that each species encodes between eight to fifteen eIF4E family members, a number surpassing that found in any other eukaryote including other alveolates (32). These analyses have shown that alveolate eIF4E cannot be grouped confidently into the groupings previously established for metazoa/plants/fungi. Instead, alveolate eIF4Es fall into three phylogenetic clades that are distinct from the three classes described for plants, fungi, and metazoans, as well as the six eIF4E family members described for trypanosomatids. The three clades can be further divided into nine sub-clades, with multiple members in each sub-clade in some species. Here we undertook a detailed examination of seven of the eight eIF4E family members in *Amphidinium carterae*, a basal core dinoflagellate. *A. carterae* represents one of the more well-studied dinoflagellate species and has begun to emerge as a “model” for dinoflagellates, (48). This species grows well in culture to high cell densities, can be grown axenically (49), is available from culture collections, has a relatively small genome of about 5.9 x Mbp per cell (11) and contains the typical dinoflagellate plastid with the peridinin pigment (50). Because of these features, we chose to work with this species to begin characterizing mechanisms of translational control using established biochemical methods with the goal of gleaning a better understanding of the complicated biology of dinoflagellates. We hope to apply these findings from a “model dinoflagellate” to the phylum as a whole.

In the current study, *A. carterae* SL-RNA was isolated and analyzed using mass spectrophotometric methods. The 5’-cap base was shown to be m^7^G. In addition, we found that the SL is very variable and is heavily modified on all bases. As measures of translation initiation factor function, we have looked at the ability of the *A. carterae* eIF4E family members to interact with cap analogues *in vitro* and *ex vivo*, their accessibility in the cytoplasmic compartment of the cell, and their ability to complement yeast conditionally deficient in eIF4E. Additionally, we have developed a novel surface plasmon resonance assay to characterize the affinities of three eIF4E family members from *A. carterae* and compared those to the well characterized murine eIF4E1A.

## Results

### The *A. carterae* spliced leader shows sequence variability

Spliced leader sequences were compiled for *A. carterae* from the Illumina RNA-seq library, Genbank SRA SRX722011, from searches using the last seven bases of the previously described 22 base spliced leader (SL), TCCGTAGCCATTTTGGCTCAAG (28). Analysis of the resulting 26,969 sequences generated is shown in the logo portrayed in Fig. 1. The relative proportions of nucleotides at each position revealed significant degeneracy in the sequence, although the previously described “canonical” sequence is the most frequent, supporting our use of that sequence for further analysis of the spliced leader 5’-cap base. Complementary analysis of Sanger sequencing data on cDNA clones from *A. carterae* confirms this degeneracy (data not shown). The first and thirteenth bases appear to be the most variable of the entire sequence with all four bases represented in near equal ratios. The eighth, nineth, fourteenth and fifteenth bases show significant variation whereas the second to seventh and the tenth to twelfth show very little. SL sequence variability is also found in nematodes and platyhelminths reflecting the existence of a large variety of SL RNA sequences (30).

**Figure 1:**
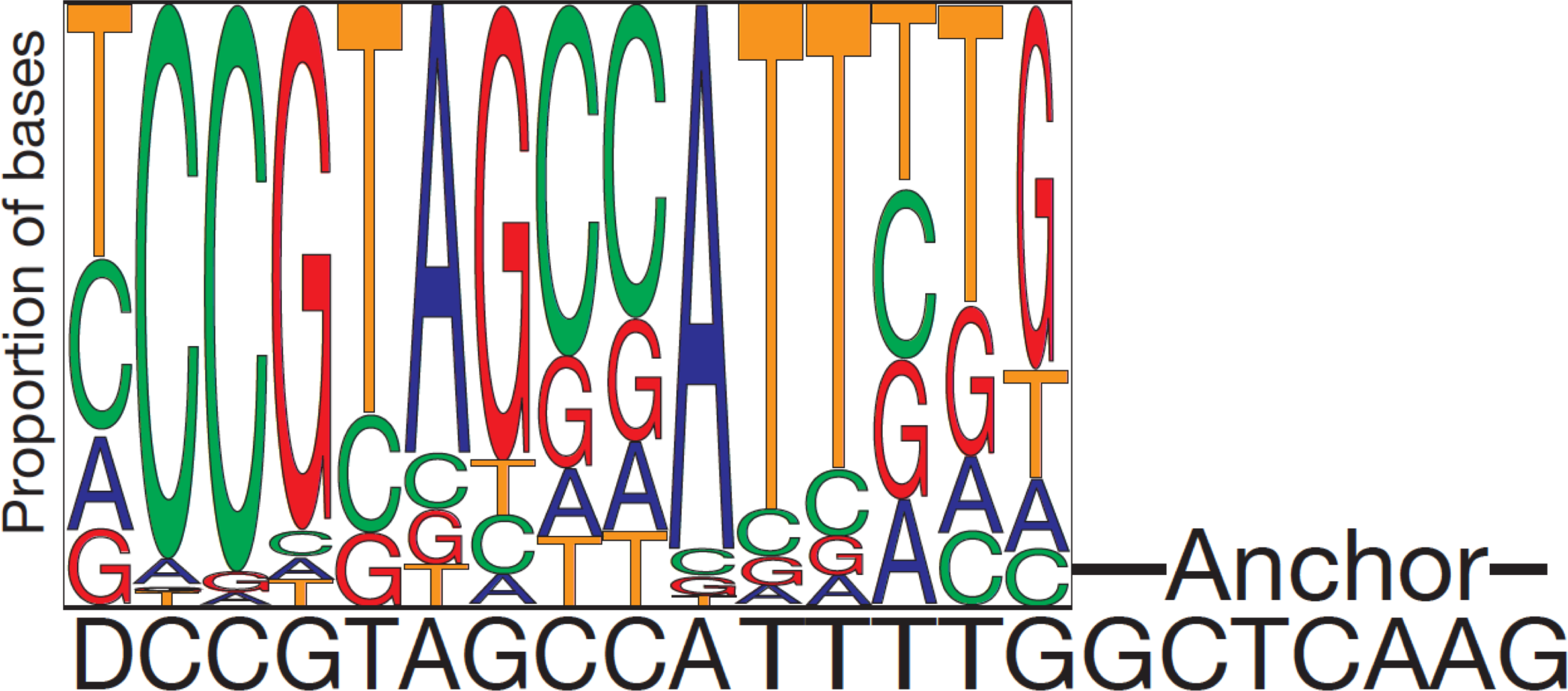
LOGO diagram of base proportions in the observed spliced leader sequences. The relative proportion of nucleotides retrieved within 50 bases of the 5-prime end of dinoflagellate transcripts is shown above the canonical spliced leader sequence from (28). The “anchor” sequence used to retrieve potential spliced leader sequences bioinformatically is shown in the right side consisting of “GCTCAAG”.

### The mRNA 5’-cap base in *A. carterae* is m^7^G and the spliced leader is hypermethylated

The identity of the 5’-cap base was of interest since other lineages that exhibit *trans-* splicing such as nematodes and trypanosomatids show unusual cap structures that interact with unusual eIF4E family members (51). We used a biotinylated oligonucleotide complementary to the canonical dinoflagellate spliced leader to recover SL RNA from a pool of small RNAs (<200 bases) isolated from *A. carterae*. To verify the efficacy of this experimental strategy, a biotinylated oligonucleotide complementary to U4 RNA was used to recover U4 RNA from the same small RNA pool.

The RNAs recovered were visualized by fractionation using an Agilent RNA chip. Both a 65-66 base SL RNA, as well as a 68 base U4 RNA were recovered (Fig. 2<u>, lanes 3 and 6</u>). The SL RNA was digested with T2 RNA nuclease producing a major product at 22 bases presumed to be the spliced leader as well as smaller bands representing methylations that inhibit further digestion (Fig. 2<u>, lane 4</u>). Such a finding may indicate incomplete digestion, despite the fact that conditions were chosen to favor full digestion. Alternatively, because RNAse T2 cannot cleave the ester bond adjacent to a 2’-O-methylated ribonucleoside (52) this could indicate that there are multiple 2’-O-methylation sites within the SL sequences that are differentially resistant to T2 nuclease. The anti-SL oligonucleotide was also used to isolate SL-containing RNA, representing mRNA, from the large RNA pool and this was digested with RNAse A. A 22-base fragment was produced confirming the size of the SL and showing that it was protected from digestion by RNAse A by duplexing with the anti-SL oligonucleotide (Fig.2<u>, lane 5</u>).

**Figure 2:**
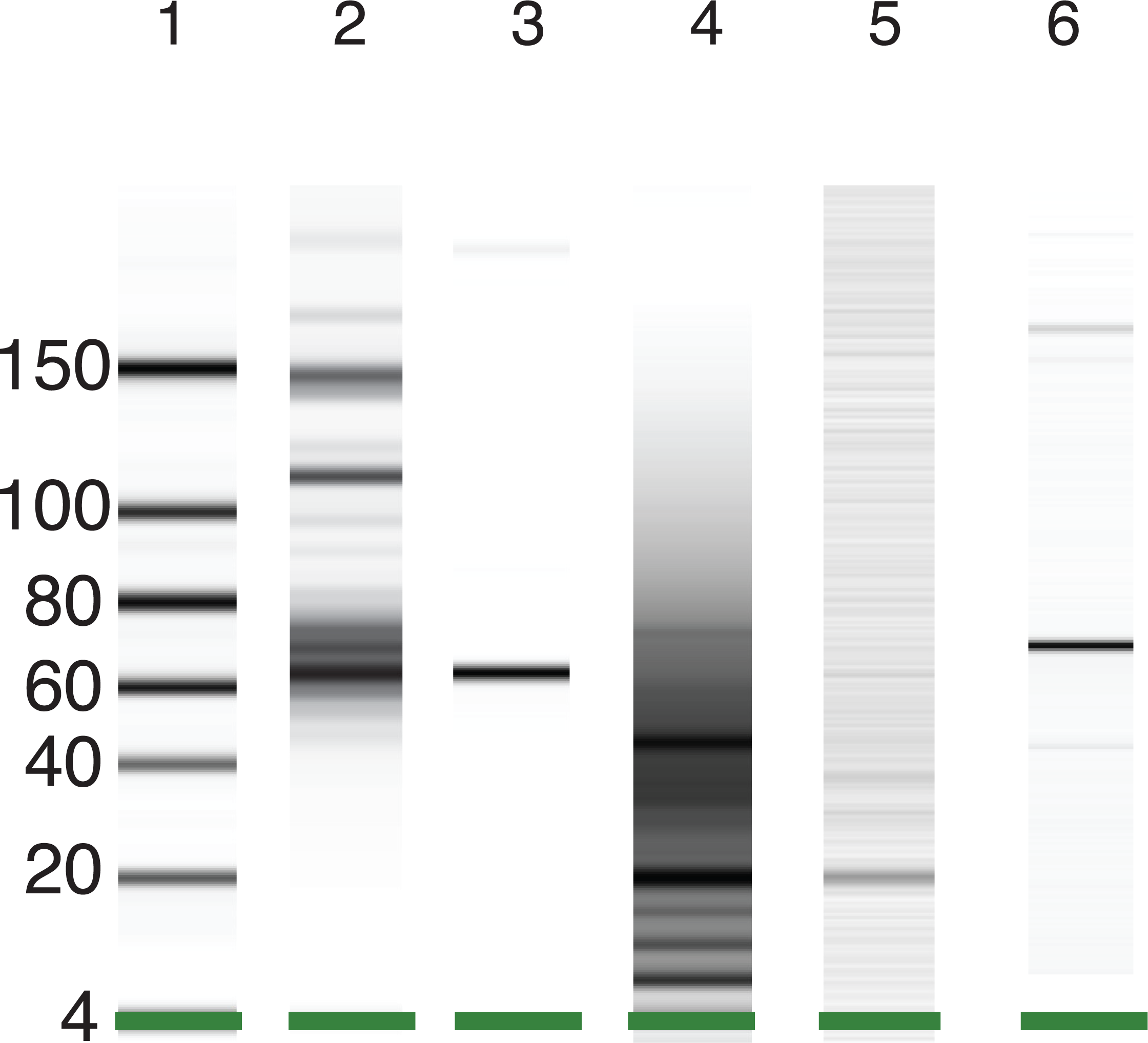
Products recovered from total small RNA using biotinylated oligonucleotides complementary to the SL and U4 RNAs. SL- and U4-containing sequences from a small RNA fraction (<200 bases) were retrieved using biotinylated oligonucleotides complementary to either SL or U4 RNA. Products retrieved with biotinylated oligonucleotides complementary to the SL were subsequently treated with T2 RNAse or RNAse A. The products were separated electrophoretically using an Agilent Bioanalyzer and compared with size standards. A virtual gel is shown here. The samples from left to right are the size standards, the total small RNA (<200 bases), the pulldown with the anti-spliced leader oligonucleotide, the T2 RNAse digest of the spliced leader pulldown, the RNAse A digest of the spliced leader pulldown, and the pulldown with the anti-U4 snRNA oligonucleotide.

Compositional analysis was performed on all of the pools including the isolated SL RNA as well as the resultant nucleotides from decapping by DCP2 (Fig. 3). The m^7^GTP nucleotide was found in both the isolated SL as well as the decapping products as expected based on an empirically determined 50 % decapping efficiency of the DCP2 enzyme established using a synthetic cap (data not shown). In contrast, m ^2,2,7^GTP was not found in either the isolated SL or the decapping. In contrast, m^2,2,7^GTP was found in the U4 pulldown, as expected, compared to a low abundance of m^7^GTP. Many other methylated products were found and, relative to the background, the 22-base SL fragment was enriched for A and G with 2’-O methylated riboses as well as m1A and m6A (Fig. 3). The 22-base SL was not enriched for uridines with ribose methylations but did contain pseudouridines. Also, in the 22-base SL compositional analysis, most if not all of the A, C and G residues were modified relative to the amount of m^7^G ideomonstrating that the dinoflagellate SL RNA is more highly modified than the cap-4 methylation pattern reported in trypanosomatid SL RNA.

**Figure 3:**
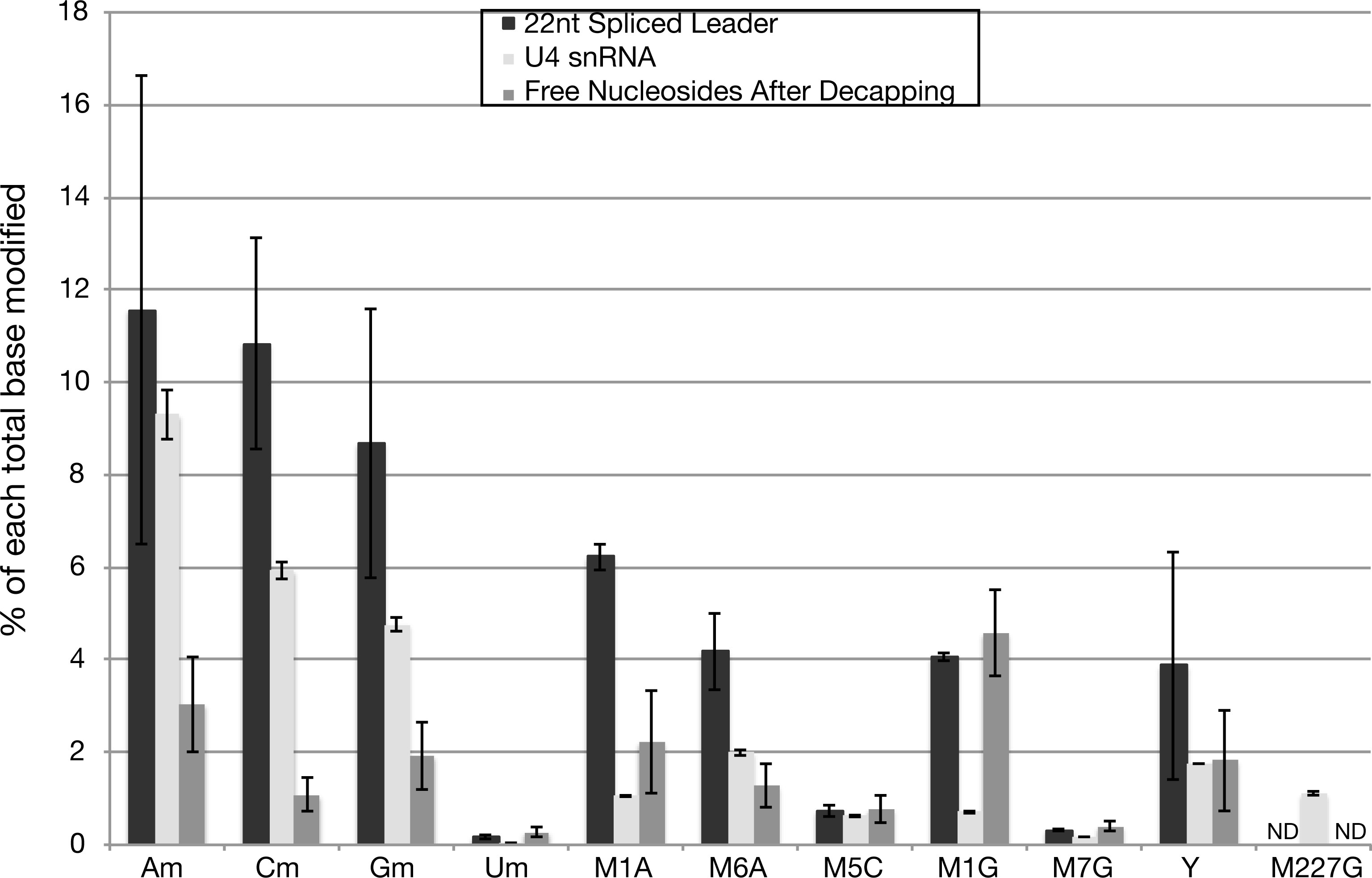
Modified bases in the 22-nucleotide SL, U4 snRNA, and decapping products. A graph of the percent of each non-standard ribonucleoside from the total detectable bases is shown with the specific moiety on the X axis and percent on the Y axis or “ND” when not detectable. The first four classes are totals for each non-standard nucleoside followed by specific moieties for which there were standards: 1-methyladenosine (M1A), 6-methyl adenosine (M6A), 5-methyl cytosine (M5C), 1-methyl guanosine (M1G), 7-methyl guanosine (M7G), pseudouridine (Y), and 2,2,7-trimethyl guanosine (M227G). The RNAse A degradation of spliced leader isolates is shown in <u>black</u>, the U4 snRNA isolate is shown in <u>light grey</u>, and the free nucleosides following decapping of the 22nt spliced leader are shown in <u>dark grey</u>. Error bars represent triplicate compositional analyses from a single sample. Moieties other than 7-methyl guanosine following decapping are likely contaminants from the total RNA pool bound to the Sepharose beads used in sequence enrichment.

### A. *carterae* capping enzymes

The 5’-cap base on mRNA or on snRNAs that donate the SL to SL trans-spliced mRNAs, is the product of three enzymatic activities: an RNA 5’-triphosphatase, a guanylyl transferase, and a methyl transferase. These are found as separate proteins in protists, or as various fusions in metazoans, fungi, or viruses (53, 54). BLAST searches were performed to identify putative capping enzymes in the *A. carterae* transcriptome. Contigs encoding the necessary domains to perform the capping reactions were identified and subsequently used to screen other available dinoflagellate transcriptomes (Table 1).

**Table 1:** Contigs from the *A. carterae* transcriptome (Genbank SRA SRX722011) presumed to play a role in capping based on annotation. Contigs from the *A. carterae* transcriptome presumed to pay a role in mRNA capping or RNA modification are listed using their contig designation (comp*****). Also listed for each contig is the expression level as fragments per thousand bases per million reads (FPKM), the eValue score and alignment length returned by BLAST, and the domain description and eValue also returned by BLAST denoting its functional role in cap structure formation.

| ***A. carterae***<br>**sequence**<br>**number** | **Guanyl**<br>**transferase**<br>**comp40464** | **Methyl**<br>**transferase**<br>**comp29574** | **Triphosphatase**<br>**Comp39307** | **O-methyl**<br>**transferase**<br>**Comp-**<br>**11589** |
| --- | --- | --- | --- | --- |
| **FPKM** | 2.8 | 2.7 | 1.9 | 13.1 |
| **eValue** | 1.00E-67 | 8.00E-40 | 3.00E-27 | 7.00E-54 |
| **Alignment**<br>**length** | 357 | 446 | 304 | 193 |
| **Domain**<br>**designation** | COG5226 | pfam032291 | Cd07470 | Pfam09445 |
| **Name of**<br>**domain** | CET1 mRNA<br>capping enzyme,<br>guanylyltransferase | Pox_MCEL<br>mRNA<br>capping<br>enzyme | CYTH mRNA<br>triphosphatase | RNA cp<br>guanine n2<br>methyl<br>transferase |
| **eValue to**<br>**domain** | 2.16E-62 | 8.41 e-29 | 1.00E-18 | 1.25E-37 |

Genes encoding all three enzymatic activities used to transfer a methylated guanosine to the 5’-end of a mRNA, or SL RNA, are present in the transcriptomes of dinoflagellates as individual open reading frames and without fusions. Contigs from the *A. carterae* transcriptome (Genbank SRA SRX722011) presumed to play a role in capping and SL modification based on annotation are shown in Table 1. Sequences encoding putative 5’-triphosphatases were common, possibly performing roles in poly-phosphate storage. The open reading frame of the *Cet1* domain common to the *beta*-subunit (RNA 5’-triphosphatase) of the *Plasmodium falciparum* 5’-RNA triphosphatase (55) was identified, as were putative methyl transferases and 2’-O-methyl transferases involved in spliced leader modifications in trypanosomes (56). A phylogenetic analysis of all three putative capping enzymes from available dinoflagellate transcriptomes along with apicomplexans was able to recapitulate the organismal phylogeny, although the methyltransferases from apicomplexans often contained repeat regions (data not shown). An additional 2’-O-methyl transferase was found that could methylate internal riboses and potentially provide more complexity in the SL structure. Additional methyltransferases that could use guanine and S-adenyl methionine as substrates were found. These were difficult to annotate but could play a role in forming the trimethyl cap of several snRNAs.

### Identification of *A. carterae* eIF4E family members and characterization based on amino acid sequence

Eight eIF4E family members were identified in our previous comprehensive phylogenetic analysis (32). Their relationships are displayed in a representative tree (Fig. 4). The eight eIF4E family members from *A. carterae* are distributed between three clades: five from clade 1; eIF4E-1a, -1b, -1c, -1d1 and -1d2: one from clade 2; eIF4E-2a: and two from clade 3; eIF4E-3a and 3b. Their characteristics are compared in Table 2.

**Figure 4.**
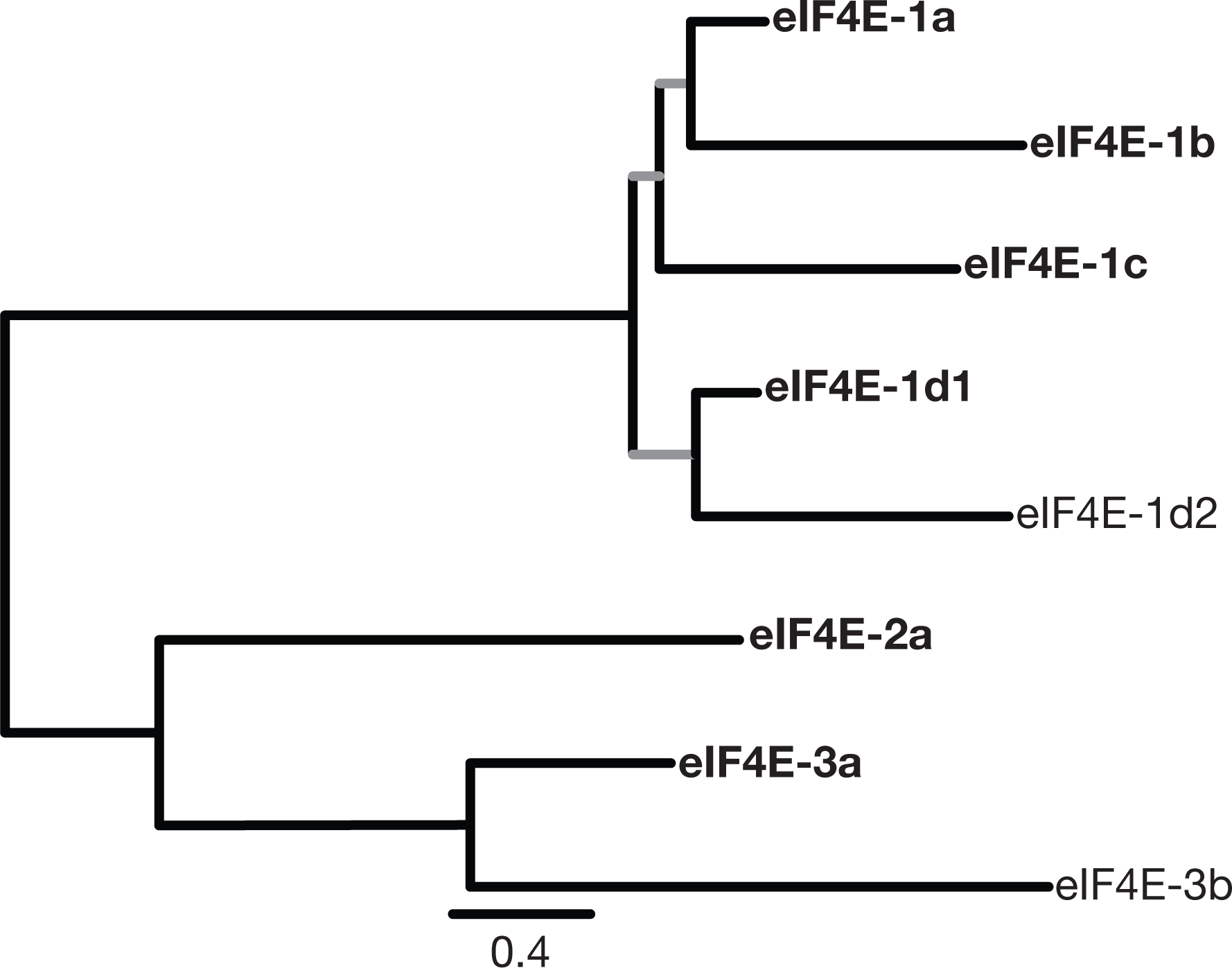
Schematic phylogeny of the eIF4E gene family in *A. carterae*, based on an amino acid alignment using maximum likelihood. The eight different eIF4E family members from this species form three major eIF4E clades in this phylogeny. Sequence labels use the nomenclature from (32). Bootstrap support is not present for the light grey branches within the eIF4E-1 clade. The eIF4E family members highlighted in bold were selected for functional analysis in this investigation.

**Table 2.**
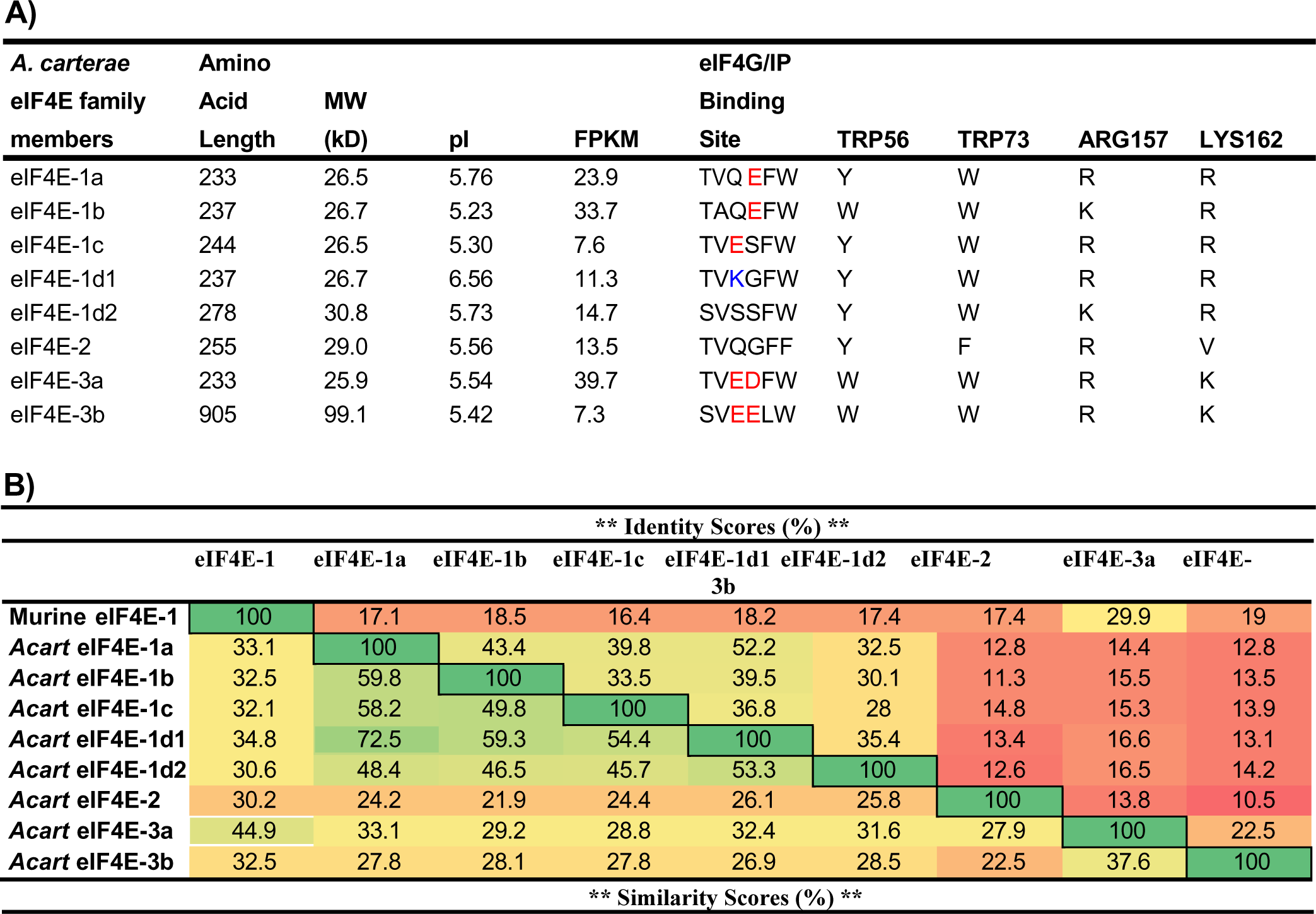
Summary of the eIF4E family members from *Amphidinium carterae*, their bio-chemical features, and their similarity to murine eIF4E-1. **A)** There are eight eIF4E family members present in *A. carterae*. They have been named according to their phylogenetic relationships as well as predictions of their biochemical functionality. We used S^35^-methionine to label recombinant proteins for our experiments. The predicted eIF4G/IP binding site is featured with red letters for negative charge and blue letters for positively charged residues. The residues, using murine numbering, are displayed for each eIF4E to highlight the differences in residues thought to be essential for cap binding. The calculated isoelectric point (pI) is an average of several alogrithms used to predict hypothetical pI values. The FPKM expression level values are taken from an Illumina HiSeqRNAseq and Trinity assembly (unpublished results). **B)** The percent identity and similarity of each eIF4E family member were compared to each other and to murine eIF4E-1. Green and yellow imply greater similarity and red implies less.

Each sequence contains a single eIF4E domain with an E value << 1E-10. Clade 1 eIF4E family members showed ≥ 28 % identity and ≥ 31 % similarity to each other, but only 30-33 % similarity to murine eIF4E-1a. eIF4E-3a had the highest similarity, 45 %, to murine eIF4E-1A (Table 2B). eIF4E-2a had the highest similarity to eIF4E-3a, at ∼28 %. The predicted amino acid sequences of seven out of the eight eIF4E family members have predicted molecular weights ranging from 25.9–30.75 kDa (from 233-278 amino acids) and predicted isoelectric points from 5.23 to 6.56 (summarized in Table 2A). The Illumina sequence data quantified as fragments per kilobase of transcript per million mapped reads (FPKM) ranged from the lowest abundance of 7.3 (eIF4E-3b) to the highest of 39.7 (eIF4E-3a), a proxy for transcript expression. The DNA sequences and translated amino acid sequences were deposited in Genbank with accession numbers listed.

Despite having low sequence identity or similarity to murine eIF4E1A (summarized in Table 2B), the core regions of each of the eIF4E family members from *A. carterae* align well with murine eIF4E1A (Fig. 5<u>, panel A</u>). A schematic highlighting the differences in residues characteristic of eIF4Es and involved in m^7^G binding or interaction with eIF4G are represented in Fig. 5<u>, panel B</u>. The eight aromatic tryptophans (or tyrosines), characteristic of an eIF4E, align with those found in murine eIF4E-1A, highlighted in yellow including Trp-56 and Trp-102 that interact with the ^7^methyl guanosine cap. eIF4E-3b provides an exception to this, with an arginine residue at the position equivalent to the Trp-102. The conserved glutamic acid at the position equivalent to E103, involved in hydrogen bonding to the m^7^G is also found in all, as are the residues involved in binding of the phosphates in m^7^G highlighted in blue, equivalent to D90 in murine eIF4E1A.

**Figure 5:**
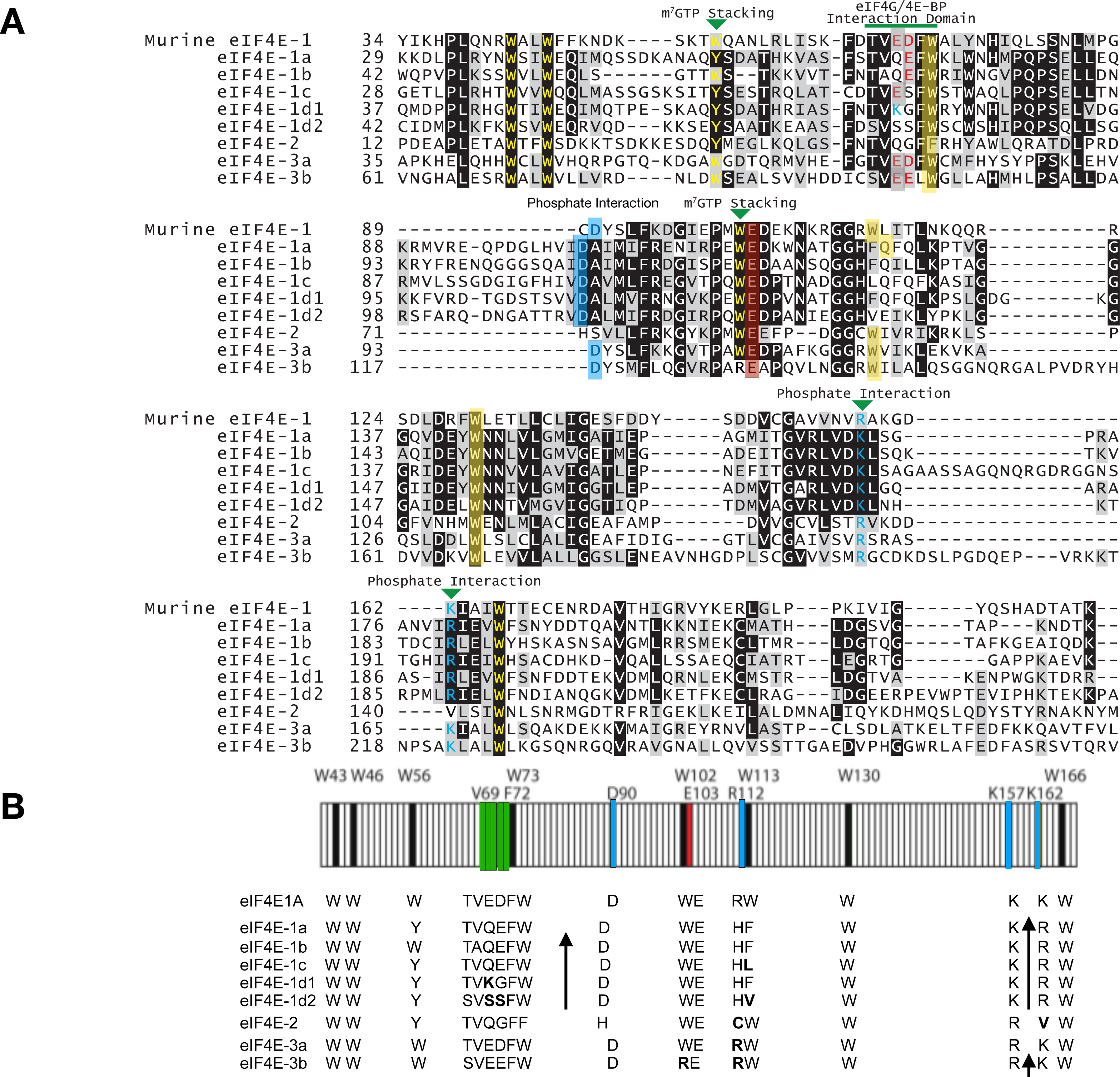
Amino acid alignment of the eight eIF4E family members from *A. carterae* with the murine class 1 eIF4E1A. **A) MUSCLE alignment** The amino acid sequences of the core eIF4E domains from the eight *A. carterae* eIF4E family members were aligned to the murine class I eIF4E, eIF4E1A, using MUSCLE. Conserved amino acids are shown in black, similar amino acids are shown in grey. The eight tryptophan residues (numbered as in the mammalian sequence) characteristic of eIF4Es are highlighted in yellow, along with the residues that are found at the corresponding positions in representative sequences from the three *A. carterae* clades. The residues known to be involved in interacting with the 7-methylguanosine cap in mammalian eIF4E1A are highlighted with a green arrowhead. The region that is associated with eIF4G/eIF4E-BP interaction in mammalian eIF4E1A is highlighted by a green bar. Residues in cyan are important for binding the phosphates of the m^7^G. In the eIF4G/4EBP interaction domain, negatively charged residues are highlighted in red and positively charged residues in blue. A. **Schematic of Important Residues:** Comparison of the conserved core region of *A. carterae* eIF4E family members with that of mammalian eIF4E1A illustrating the important binding regions for the m^7^G cap and eIF4G. The eight conserved tryptophan residues (numbered as in the mammalian sequence) are shown in black along with the residues that are found at the corresponding positions in representative sequences from the three *A. carterae* clades. Residues in yellow are important for binding the phosphates of the m^7^G (the aspartate at position 90 coordinates binding by arginine157), while the eIF4G binding motif (S/TVxxFW) is shown in blue. The glutamate at position 103 (red) is involved in hydrogen bonding to the m^7^G. Note that dinoflagellate clade 1 eIF4Es have an insertion (↑) of 12–13 amino acids between positions equivalent to W73 and W102, as well as an insertion of 7–9 amino acids between W130 and W166.

Immediately apparent are two amino acid insertions in all clade 1 eIF4Es between the position equivalent to Trp-73 and Trp-102 in murine eIF4E1A, a feature not seen in any plant or metazoan eIF4E family member but found in all dinoflagellate species (32). Another key difference is in eIF4E-2 in which an uncharged valine is present at the position equivalent to Lys/Arg-162 in murine eIF4E-1A and present in all other dinoflagellate eIF4E family members (32). There are also differences in the eIF4G binding region. Like murine eIF4E1A, *A. carterae* eIF4E-1a, -1b, -1c and eIF4E-3a and -3b contain negatively charged amino acids in this region, while eIF4E-1d1 is unique in that it contains a positively charged lysine in this region. Both eIF4E-1d2 and eIF4E-2 contain only uncharged residues in this region.

eIF4E-3b is an outlier with a molecular weight of 99.1 kDa (905 amino acids), although its isoelectric point, 5.42, is within the range of all other eIF4E members from *A. carterae*. eIF4E-3b additional domains; a predicted “2OG-Fe(II) oxygenase superfamily” domain (pfam13532) with an E-value of 1.12E-19 and an “alkylated DNA repair protein” domain (COG3145) with an E-value of 5.57E-11 at the amino-terminus (Fig.5<u>-Suppl. Table 1</u>). This is unlikely to be a sequencing artifact since such a form of eIF4E has also been described for six additional dinoflagellate species and represents the only 2OG-Fe(II) oxygenase:eIF4E fusion described from any available eukaryotic sequences (32).

### Structural prediction and modeling of *A. carterae* eIF4E family members

Amino acid sequence from each eIF4E family member was submitted to the Phyre2 Protein Fold Recognition Server. The resulting predicted structure was output as a PDB file that was then used to model the protein structure using the Visual Molecular Dynamics software (Fig.5<u>-Suppl. Fig.1</u>). Due to the evolutionary distance and amino acid differences from the eIF4E crystal structures, the cap-stacking residues do not always fold into the expected positions to interact with the cap structure. However, the positively charged residues interacting with the phosphate bridge usually fall into place near the phosphate bridge. eIF4E-2 is notable for its uncharged valine residue in place of a positively charged lysine in the position required for phosphate bridge interaction.

Each of the eIF4E family members model well to murine eIF4E1A structures (Fig. 5<u>-Supp. Fig. 1</u>). Each eIF4E from *A. carterae*, except eIF4E-3b, was modeled with high confidence (100 %) across 51-87 % of the sequence to *H. sapiens* or *M. musculus* eIF4E1A using Phyre2 Protein Fold Recognition algorithm. The output PDB file was used to visualize the residues expected to interact with and bind the mRNA cap structure (Fig. 5<u>-Supp. Fig. 1</u>), highlighted in Fig. 5<u>, panels A and B</u>. For each sequence, the core structure resembles the cup-shape associated with an eIF4E translational initiation factor. The two tryptophans that sandwich the 7-methylguanosine do not always stack appropriately in the predicted structures. However, the two positively charged residues associated with interaction of the negatively charged phosphates of the cap structure model into appropriate positions for making such contacts, except in eIF4E-2, in which Lys-162 has been replaced with a hydrophobic valine, Val-162. eIF4E-3b modeled with >90 % confidence for 47 % of the residues to *H. sapiens* eIF4E-1 (not shown). The two domains of eIF4E-3b modeled separately to two protein families, but together form a single multi-domain protein (not shown). The top hit for the eIF4E domain of eIF4E-3b had 30 % identity and modeled with 100 % confidence, while the 2OG-Fe(II) oxygenase domain of eIF4E-3b had 27 % identity with an alpha-ketoglutarate-dependent dioxygenase (PFAM: PF13532) and modeled with 99.7 % confidence. The characteristics of eIF4E-3b have not been investigated in this study due to its lack of conservation in the phylogenetic analysis amongst dinoflagellates and the additional domain present. It is still of interest since the alpha-ketoglutarate-dependent dioxygenase (AlkD) domain may have a role in nucleotide methylation (57).

### Transcript levels of *A. carterae* eIF4E family members

Transcript levels of all eight eIF4E family members, expressed as qPCR cycle thresholds, were determined at the mid-day and mid-night time points (primer sequences shown in Fig. 6<u>-Supp.l Table 1</u>). The transcript levels varied coordinately between mid-day to mid-night (Fig. 6). Transcript levels were somewhat higher for all eIF4E family members at the mid-night point. eIF4E-3a is the most highly expressed at the transcript relative to the other eIF4E family members while eIF4E-3b is the lowest at mid-day or mid-night, consistent with the FPKM values. The expression of other genes, such as rpl7, psbO, psbB, and the ribosomal RNA gene LSU were found to display the same differences in expression from mid-day to mid-night (not shown).

**Figure 6:**
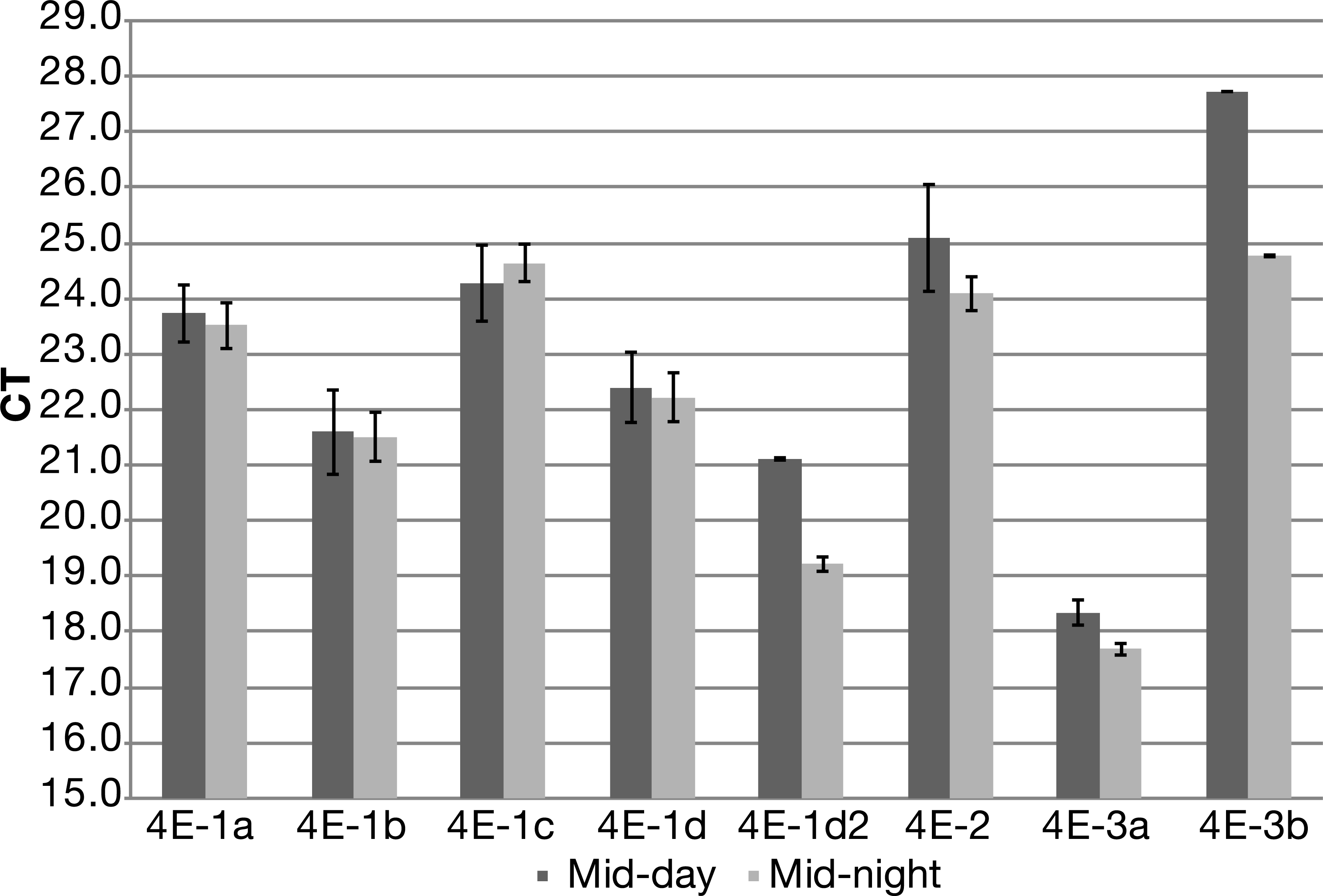
Quantitative-PCR cycle thresholds of each eIF4E family member at a mid-day and mid-night time point on a diel cycle. The relative transcript abundance of each eIF4E family member is shown as a cycle threshold for the mid-day and mid-night time points of *A. carterae* cultures maintained on a 14:10 light:dark cycle.10 ng RNA was reverse-transcribed using random primers and used as template for qPCR, as outlined in <u>Materials and Methods</u>. cDNA was measured using SYBR green as an indicator.

### Quantification of eIF4E protein levels from *A. carterae*

eIF4E protein levels were determined by western blot analysis using affinity antibodies generated by GenScript Inc, Washington, DC. The peptide sequences used to generate the antibodies are shown in Fig. 7<u>-Suppl. Table 2</u>. The specificity of antibodies used for western analysis is shown in Fig. 7<u>-Suppl. Fig 1</u>. In contrast to transcript abundance, eIF4E-1a is the most highly expressed of the family members at the protein level, present at approximately 2.34×10^6^ molecules per cell, about 5.5-fold greater than the next most abundant member, eIF4E-1d1, at 4.28×10^5^ molecules per cell (Fig. 7).

**Figure 7:**
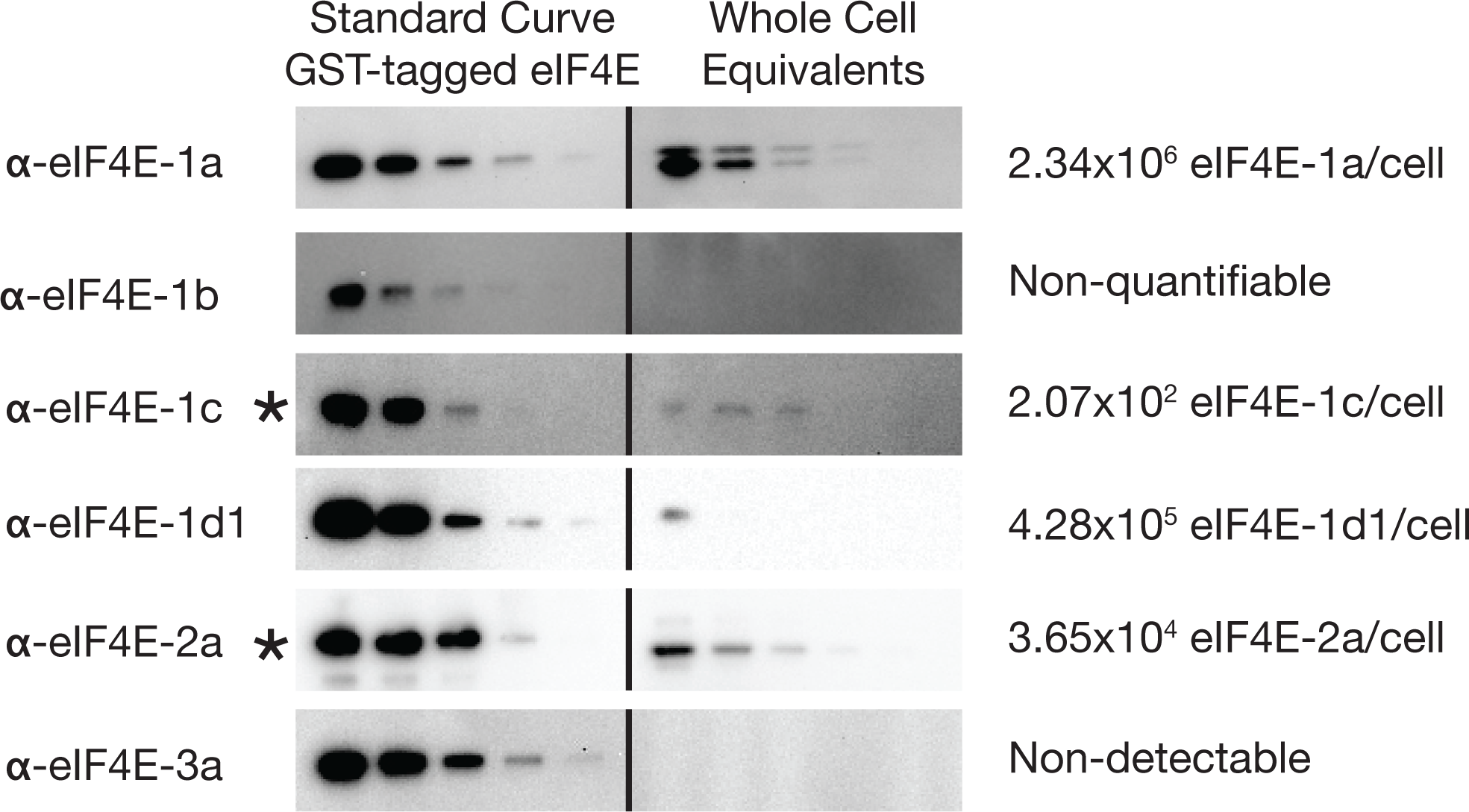
Quantification of protein abundance of each eIF4E family member in *A. carterae*. Recombinant GST-tagged protein was purified from either the soluble or insoluble fraction of an *E. coli* lysate. Recombinant protein was quantified and diluted into a two-fold dilution standard curve starting at 25 nanograms for eIF4E-1a and eIF4E-1d1 or 3 ng for eIF4E-1b, eIF4E-1c, eIF4E-2, and eIF4E-3a. This was used to compare to a dilution curve of *A. carterae* suspended and boiled directly in SDS-PAGE samples buffer. Four two-fold dilutions were loaded per well starting at 250,000 cell equivalents for eIF4E-1a and eIF4E-1d1 or 500,000 cell equivalents for eIF4E-1b, eIF4E-1c and eIF4E-2. The pixel densities of each band from the standard curve were used to calculate the nanogram quantities and converted into molecules per cell based on the molecular weight and cell equivalents loaded in that well.

The amount of recombinant protein used in the standard curve had to be lowered by 10-fold for eIF4E -1c (2.07×10^2^ molecules per cell) and eIF4E-2a (3.65×10^4^ molecules per cell) in order to be in a suitable range. In whole cell lysates, eIF4E-1b was detected when the blot was overexposed without recombinant protein present (data not shown), but could not be detected in the dynamic range for quantification. However, eIF4E-3a was not detectable from lysates using western blot, even with overexposure. An additional eIF4E-3a antibody was generated using a second epitope and used in the same analysis. This antibody also failed to detect expression of eIF4E-3a in the dynamic range for quantification (data not shown).

To complement the western blot analyses, we performed a small-scale proteomics analysis of proteins from *A. carterae* in the 27-33 kDa range isolated from whole cell lysates and fractionated by SDS-PAGE. Significant coverage (greater than 3 independent peptides mapped) was shown for peptides mapping to eIF4E-1a, -1b, and -1d1 (Fig.7<u>-Suppl. Fig. 2</u>). Despite the difficulty in measuring the expression of eIF4E-1b by western blot analysis, of cell lysates, eIF4E-1b peptides were mapped in the proteomics analysis. Peptides mapping to eIF4E-3a were not found, consistent with the data from western analysis, again indicating its lack of expression at the protein level. These data provide confidence that eIF4E-1a, -1b, and -1d1 are translated to protein and found within the predicted size (27-33 kDa) (Table 2A). eIF4E-3b (at 99.1 kDa), would not have been found in this size range analyzed. However, eIF4E-3b was not observed in a proteomic study in *Polarella* which did not rely on size fractionation consistent with our results(14). A similar complement of Clade 1 eIF4Es was observed in *Symbiodinium* species in a comparative genomics study using more inclusive methods (58).

### Recombinant eIF4E-1 and eIF4E-3 family members from *A. carterae* interact with the m^7^GTP cap analogue, but eIF4E-2 does not

To test the ability of the eIF4E family members from *A. carterae* to function as cap-binding proteins, we assayed the ability of the recombinant proteins to bind to cap analogue. Since the 5’-base was found to be m^7^GTP, the cap binding ability of *A. carterae* eIF4E family members was initially assessed using m^7^GTP-Sepharose beads (Fig. 8 and Fig. 8<u>-Supp. Fig. 1</u>). Recombinant eIF4E proteins were transcribed/translated *in vitro* from codon-optimized pCITE4a constructs and radiolabeled using ^35^S-methionine, essentially as described previously (59). The translated products were incubated with an m^7^GTP-Sepharose bead slurry, and the unbound, and bound fractions were analyzed by direct counting of bound and unbound fractions, as well as by SDS-PAGE and autoradiography. *Renilla* luciferase was used as a negative control. eIF4E-1a, -1b, -1d1, and eIF4E-3a bound m^7^GTP-Sepharose, as did the positive control, *C. elegans* IFE-1 (60). Neither eIF4E-1c nor eIF4E-2a bound well to m^7^GTP in this assay (Fig. 8 and Fig. 8-<u>Supp. Fig. 1</u>). Binding to trimethylated-guanosine (TMG), the cap structure present in *C. elegans trans*-spliced mRNAs, was also tested using TMG-Sepharose (a generous gift of Edward Darzynkiewicz, Ph.D., University of Warsaw). eIF4E-1a, eIF4E-1b, eIF4E-2 and eIF4E-3a did not bind the TMG cap analogue, although eIF4E-1c and eIF4E-1d1 showed some affinity for the TMG cap analogue. The known TMG-binding eIF4E, IFE-1, of *C. elegans* (NCBI Gene ID: 176755), bound well in both assays (60), while the *Renilla* luciferase came out in the unbound fraction.

**Figure 8:**
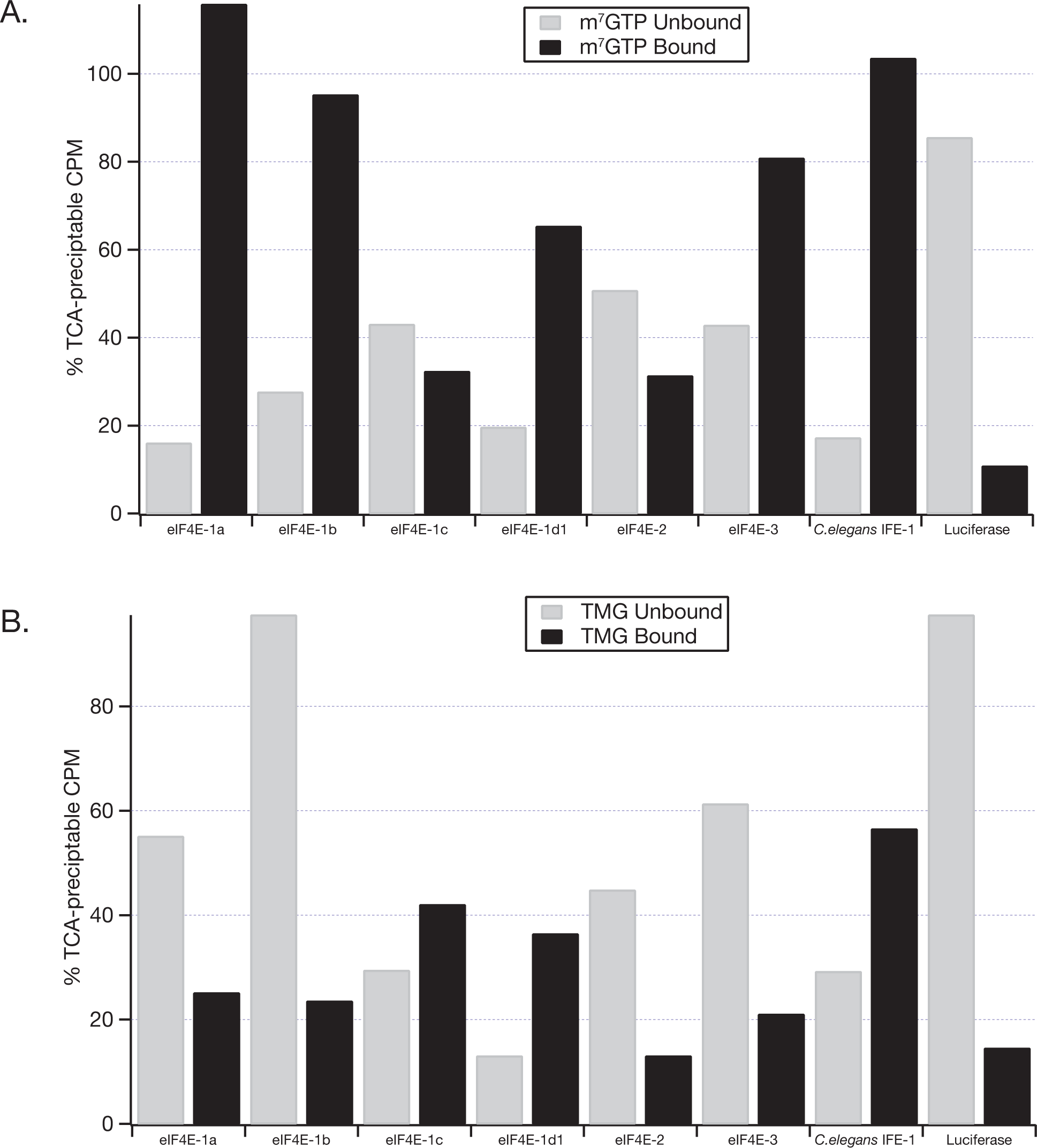
Assessment of the m^7^GTP and TMG binding capability of eIF4E family members from *A. carterae*. S35-methionine-labelled eIF4E family members were produced using a rabbit reticulocyte *in vitro* expression system. The ability to interact with m^7^GTP vs TMG cap was assessed by measuring the CPM of the unbound and bound fraction after loading, equilibrating, and washing the cap-Sepharose column. The counts were expressed as a percent of the total incorporation into each family member. The *A. carterae* eIF4E family members were compared to a positive control, *C. elegans* IFE-1, and a negative control, luciferase.

### Only eIF4E-1a can be retrieved from *A. carterae* cell lysates by binding to m^7^GTP-Sepharose beads

To examine which endogenous eIF4E(s) from *A. carterae* have the ability to act as a cap binding translation initiation factor, we tested the ability of native protein derived from a cell lysate to bind to m^7^GTP. Using m^7^GTP-Sepharose beads, only eIF4E-1a can be isolated from *A. carterae* cell lysates on western blots (Fig. 9). Other family members, eIF4E-1b, eIF4E-1c, or eIF4E-2a, were found in the unbound fraction, or in the case of eIF4E-1d, only in the pelleted insoluble fraction generated after cell lysis. eIF4E-3a was not found in any fraction. These data demonstrate that eIF4E-1a is soluble and accessible in the cell lysate for binding to the cap analogue beads, but that eIF4E-1b, -1c, and 1d1, and -2a may be compartmentalized or unable to bind the cap structure due to modifications or binding to other partners.

**Figure 9:**
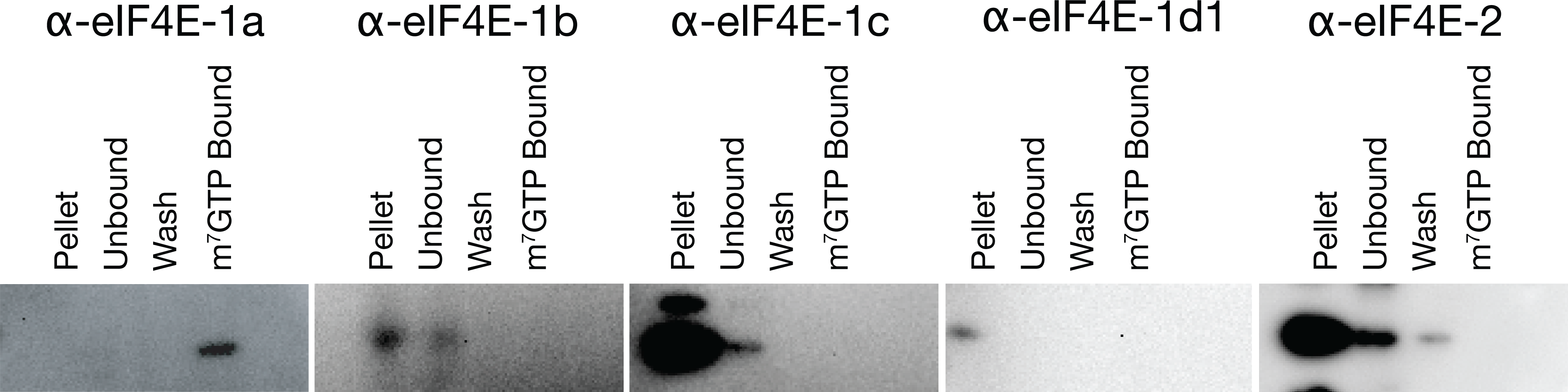
Only eIF4E-1a can be retrieved by m^7^GTP-Sepharose chromatography from cell lysates of *A. carterae*. Cell lysate was generated from a mid-day actively growing culture of *A. carterae*. Proteins bound to m^7^GTP-Sepharose were analyzed by western blot using antibody specific for each eIF4E family member. eIF4E-3a was not included in this analysis since we could not find expression of it using western blot with two separate antibodies.

### *A. carterae* eIF4E-1a and -1d are functionally equivalent to human eIF4E-1 in *S. cerevisiae*

Although there is considerable sequence divergence between human eIF4E-1 and *S. cerevisiae* eIF4E (31 % identity), the mammalian factor can sustain growth of yeast conditionally deficient in eIF4E. The yeast strain, JOS003 (61), previously developed to compare the functionality of zebrafish eIF4E-1A and eIF4E-1B (59) was used to determine the functionality of *A. carterae* eIF4E family members. The JOS003 strain lacks the endogenous yeast eIF4E gene and expresses human eIF4E-1 inserted in the pRS415 leu (-) vector under the control of the galactose-dependent and glucose-repressible GAL1 promoter. As a consequence, strain JOS003 is able to survive in medium containing galactose as carbon source, but is not viable in medium containing glucose due to depletion of the human eIF4E-1. Growth of JOS003 in glucose can be mediated by ectopic expression of a functional eIF4E, regulated by a promoter in the pRS416 ura (-) vector, which is active in the presence of glucose. Following transfection and selection on media lacking uracil, the yeast cells containing control vector, or constructs for the expression of eIF4E-1a, -1b, -1c, 1d1, eIF4E-2a and eIF4E-3, were streaked on selective plates; SD - Ura, -Leu containing either galactose or glucose as carbon source (Fig. 10).

**Figure 10.**
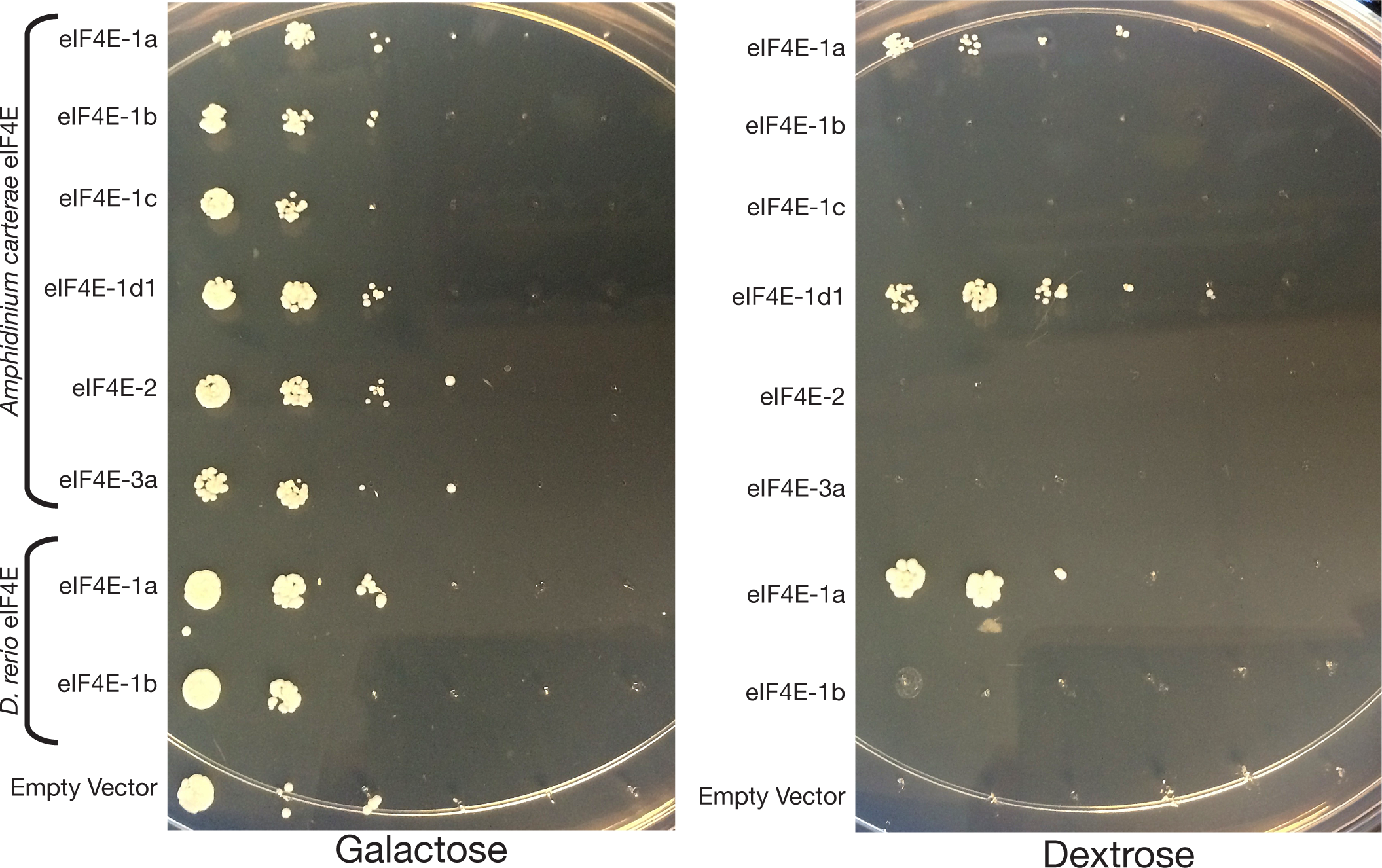
*A. carterae* eIF4E-1a and eIF4E-1d complement yeast lacking the endogenous eIF4E gene. The *S. cerevisiae strain*, JOS003 (61), was transformed with the Ura-selectable vector, pRS416GPD, that was either empty or contained cDNAs encoding one of the following: *A. carterae* eIF4E-1a, -1b, -1c, -1d1, eIF4E-2a, eIF4E-3a. pRS416GPD constructs containing zebrafish eIF4E-1A and -1B were also included as positive and negative controls, respectively (59). Following selection on SC medium with galactose but lacking uracil and leucine, yeast from the resulting single colonies were transferred to YP-agar media containing G418 and either glucose (right), or galactose (left) as carbon source and allowed to grow at 30 °C for 72 h.

Zebrafish eIF4E-1A and eIF4E-1B were used as positive and negative controls, respectively. As previously reported, zebrafish eIF4E-1A was capable of complementation, while eIF4E-1B was not, reflecting its inability to interact with the cap structure or eIF4G (59). Of the *A. carterae* eIF4Es, only eIF4E-1a and eIF4E-1d1 were able to rescue growth of this yeast test strain (Fig.10). Of note, the eIF4E family member with the highest identity/similarity to *S. cerevisiae* eIF4E and human eIF4E-1A, *A. carterae* eIF4E-3a, was not able to rescue growth. Expression of the *A. carterae* proteins was confirmed by western analysis (Fig. 10<u>-Supp. Fig. 1</u>). Interestingly, eIF4E-1d2 did not complement, despite being the closest related family member to eIF4E-1d1, another reason this eIF4E family was not pursued for further study (Fig. 10).

Since codon optimization may have a role in protein synthesis efficiency and stability in a heterologous host (62, 63) we generated codon-optimized translation constructs for the *A. carterae* eIF4E family members. In general, eIF4Es from phylogenetically distant organisms are considered too divergent to complement a missing yeast eIF4E. For instance, although *Drosophila* eIF4E-1, -2, -3, -4 and eIF4E-7 (all Class I eIF4E family members) are able to complement the lack of eIF4E in a yeast CDC33 (eIF4E)-knockout strain (39), eIF4Es from the phylogenetically more distant trypanosomes cannot (64). In contrast, codon optimization did not have any additional positive effect on the ability of eIF4E-1a to rescue growth; only *A. carterae* eIF4E-1a and eIF4E-1d1 complement the *S. cerevisiae* eIF4E conditional knockout (Fig. 10<u>-Supp. Fig. 2</u>).

### Analysis of cap-binding by surface plasmon resonance

We used binding to bead-bound cap analogs as a first approximation of cap binding functionality. However, such assays provide no quantitative assessment of cap binding ability. To obtain equilibrium affinity constants for interactions between *A. carterae* eIF4Es and the mRNA cap we used Biacore surface plasmon resonance technology. Surface plasmon resonance was used because previous work from other groups have established this as a viable assay for measuring the affinity of eIF4E for cap analogs (65, 66). In addition, the version of the SPR technology we used, Biacore T-200, is able to detect very small changes in response units, and therefore provides a direct measure of mass changes caused by the low molecular weight cap structure, with low background (personal communication, Michael Murphy, Ph.D., Biacore). We used two of the *A. carterae* eIF4E family members likely to be involved in general translation according to the bead binding data; eIF4E-1a and eIF4E-1d1. In addition, we tested one eIF4E that is predicted to be missing the necessary electrostatic interactions to interact with the cap structure; eIF4E-2a. Our strategy was to use recombinant glutathione-s-transferase (GST)-tagged eIF4E bound to an anti-GST chip with the different cap analogues injected over the chip. GST-tagged proteins were purified by glutathione affinity column chromatography, dithiothreitol treatment to disperse inactive (or multimeric) complexes and gel filtration to isolate GST-fusion protein dimers of the appropriate molecular weight (Table 3<u>-Supp. Fig. 1</u>). Our measure of protein folding was based on the ability to purify protein using GST-affinity chromatography, elution of a single peak with high symmetry from the gel filtration column, and elution at a volume calculated to correspond to a molecular weight that approximated the expected molecular weight for a GST-fusion dimer compared to a protein standard curve (67–69).

**Table 3:** The binding affinity of purified *A. carterae* eIF4E family members for three separate cap structure analogs using SPR. Recombinant GST-tagged eIF4E proteins were purified from *E. coli* and loaded onto a surface plasmon resonance chip equipped with an anti-GST antibody. Binding experiments were carried out in phosphate-buffered potassium buffer (20 mM sodium phosphate pH 7.5, 150 mM KCl, 0.05% Tween-20) at 25°C. A two-fold dilution series (250-1.95 μM for the dinucleotide and 62.5 - 0.97 μM for the mononucleotide) was used to measure the change in response for each cap structure analogue. All dissociation constants (KD) were calculated based on a dose response curve *except for the affinity of eIF4E-1d1 and mouse eIF4E-1 for m^7^GpppC which was calculated based on kinetic curve fit by Biacore software algorithms.

| eIF4E | m<sup>7</sup>GpppG<br>K<sub>d</sub> (μM) | m<sup>7</sup>Gppp<br>K<sub>d</sub> (μM) | m<sup>7</sup>GpppC<br>K<sub>d</sub> (μM) | GpppG<br>K<sub>d</sub> (μM) |
| --- | --- | --- | --- | --- |
| eIF4E-1a | 17.5 +/-4.68 | 3.47 ± 1.24 | 10.8 ± 3.1 | 1726+/-3700 |
| eIF4E-1d1 | 34.5+/-18.6 | 3.27 ± 0.78 | 82.4* | 1006 ± 110 |
| eIF4E-2 | 468.8+/-220 | No affinity | No affinity | 665.3 ± 19.0 |
| eIF4E-1<br>(mouse) | 59.7+/-49.3 | 0.23 ± 0.56 | 69.8* | 316.4 ± 38.0 |

We tested the affinity of each fusion protein against the cap analogs: m^7^GpppG, m^7^GpppC, m^7^Gppp, GpppG, and Gppp. Steady state affinities were calculated from a dose response curve fit of Rmax changes with each successively higher concentration of analyte (cap analog) (Table 3<u>-Suppl. Figs 2, 3, & 4</u>). We verified our methodology using mouse eIF4E-1A for which cap binding affinity has been defined (67–69). For most of the cap analogs tested, the fast changes in response units (RUs) made it impossible to calculate a kinetic fit to the sensorgrams. Instead, only steady state affinity could be derived by plotting the response changes (a single time point of data along the 60 sec injection) against the concentration of analyte in that injection. A kinetic fit of the sensorgram was only possible with eIF4E-1d1 and murine eIF4E-1A when tested with m^7^GpppC. A fit of the curve produced reliable data for the “on” and “off” rates with a calculated Kd affinity based on the K_on_ and K_off_ rates.

Using our procedure, murine eIF4E-1A has a Kd of 0.2 μM for m^7^GTP, within the range of other studies, (68, 69). eIF4E-1a and -1d from *A. carterae* have more than a ten-fold lower affinity for m^7^GTP (3.5 and 3.3 μM) than murine eIF4E1A. However, for m^7^GpppG, both eIF4E-1a and -1d (Kd of 18 μM and 35 μM, respectively) have approximately the same affinity, within the margin of error, as murine eIF4E1A (Kd = 60 μM). Furthermore, *A. carterae* eIF4E-1a has a higher affinity than murine eIF4E1A for m^7^GpppC (11 μM versus 70 μM) (Table 3).

In contrast, eIF4E-2a did not show any response for the cap structures m^7^GTP and m^7^GpppC, and very low affinity for m^7^GpppG (Kd = 470 μM), in line with functional predictions. No eIF4E displayed any affinity for GTP (data not shown) and only displayed very low affinity for the GpppG non-methylated dinucleotide (Table 3). The kinetics of binding were different for the m^7^GpppC cap structure, with a very slow “on” (92.2 – 87.4 M-1 S-1) rate and a relatively slow, but measurable “off” (0.0076 – 0.006 S-1) rate for eIF4E-1d1 and murine eIF4E1A, but not for *A. carterae* eIF4E-1a (Table 3<u>-Suppl Figs. 2, 3 & 4</u>).

## Discussion

Enormous genetic content, little transcriptional regulation, and extreme species diversity render dinoflagellates an interesting, but challenging, lineage to explore. Dinoflagellates rely heavily on post-transcriptional control for the regulation of gene expression, showing little transcriptional regulation, but seem to rely on post-transcriptional controls, we looked for innovations in mRNA recruitment unique to dinoflagellates that might enable specific translational control. We focused on translation initiation looking at the mRNA 5’-cap base structure, the spliced leader sequence, as well as the unique set of eIF4E family members. Most of what we know of the initiation of protein synthesis comes from yeast, mouse, and human cell lines, phylogenetically distant from dinoflagellates (at least 700 Ma). However, draft short-read genome assemblies (14, 33, 34, 70), as well as accumulating transcriptome data (5) have allowed us to begin to build a picture of the dinoflagellate translational toolbox (32).

### Significance of SL diversity and 5’-cap base in *A. carterae*

We have determined that the 5’-cap base of the *A. carterae* SL RNA is m^7^G, as is found in the majority of non-SL mRNAs across eukaryotes and the SL sequences of trypanosomatids. In trypanosomatids the next four nucleotides in the SL are methylated to give m^7^Gppp_3_^6,6,2^Apm^2’^Apm^2’^Cpm_2_^3,2’^U referred to as a cap-4 structure (71). This is in contrast to the 5’-cap base found in nematode SL sequence which is 2,2,7-trimethyl guanosine characteristic of most small nuclear RNAs (72).

Spliced leader (SL) *trans*-splicing is mediated by the spliceosome and allows the replacement of 5’-end of pre-mRNA by the 5’(SL)-end of SL-RNA co-equipped with a methylated 5’-cap structure. Spliced leader *trans*-splicing was first observed in trypanosomatids (73), although it has been observed in many phylogenetically unrelated eukaryotes including euglenozoans, perkinsozoans, dinoflagellates, along with animal branches such as rotifers (74), cnidarians (75), nematodes (76), platyhelminthes (77), and chordates (78). Bioinformatic comparisons of SL-RNAs from various eukaryotic taxonomic groups have revealed similarities in secondary structures of most SL-RNAs and a relative conservation of their splice sites (SSs) and Sm-binding sites (30). Since such structural and functional similarities of SL-RNAs are unlikely to have evolved repeatedly many times, it seems likely that SL trans-splicing had an early origin, but that multiple losses occurred independently in most eukaryotic lineages (31)

Consistent with the ∼60 bp observed in other dinoflagellates (79), the SL RNA in *A. carterae* is estimated to be 65 bp (Fig. 2). Since the original discovery of SL *trans*-splicing in dinoflagellates (15, 28), a variety of sequences and genomic arrangements have been found (79). The SL RNA genes in dinoflagellates are found in tandem repeats and also as SL-5S gene clusters with the ∼60 bp SL RNA being largely conserved but showing variability (14, 79). It is therefore not surprising that we find variability in the observed spliced leader sequences from *A. carterae* transcripts (Fig. 1). Such variability is consistent with that found in Euglenozoa, Nematoda and Platyhelmintha (30).

Dinoflagellates utilize *trans*-splicing, as well as encoding a large number of eIF4E family members. m^7^G is the 5’-cap base, followed by SLs with modified bases. We have shown that the sequence heterogeneity within the SL is noteworthy in *A. carterae*. As in other lineages, it is possible that different SL sequences recognize different subsets of mRNAs (reviewed (30). It is not known whether SL sequences change through the diel cycle or with different stressors. From our transcriptomic data sets, the first nucleotide and thirteenth nucleotides of *A. carterae* SL show almost equal representation by all four bases, the eighth, nineth, fourteenth and fifteenth bases show significant variation, whereas the second to seventh and the tenth to twelfth show very little. It has been shown that mRNAs that differ in their cap-proximal nucleotides can display differential affinity towards eIF4E, as is observed in mouse embryonic fibroblasts in which differential translation of transcript isoforms is part of the stress response (80). This provides one possible mechanism for how SL and eIF4E can regulate gene expression together.

The dinoflagellate SL appears to have a modified base structure that somewhat resembles that of the trypanosome cap-4 structure, m^7^Gpppm_3_^6,6,2’^Apm^2’^Apm^2’^-Cpm_2_ ^3,2’^U, cap-4 (81), but with modifications found on all bases. It should be noted that the extent of modification of bases in the complete SL of trypanosomes has not been determined only that *Trypanosoma brucei*, a minimal level of mRNA cap ribose methylation has been shown to be essential for viability (82), and hypermethylation of the trypanosome cap-4 maximizes translation rates in *T. brucei* (82). Although the SL of trypanosomatids is referred to as a cap-4 structure, it is not clear what contributions to eIF4E binding the 5′-modified SL structure plays, leading to the philosophical question, what can be considered the contribution of a 5’-cap base and what should be considered the function of the SL? The data presented here allude to an overall complexity in the *A. carterae* SL that may be greater than the cap-4 structure of trypanosomes.

RNA modifications alter the properties of the four nucleotides, thereby influencing the inter- and intra-molecular interactions of the RNAs that carry them (83). Methylation of riboses at 2’-OH group is one of the most common RNA modifications found in number of cellular RNAs, as is modification of adenosine to N6-methyladenosine (m^6^A), and the isomerization of uridine to pseudouridine (83). 2′-O-methylation typically stabilizes RNA helices by increasing base-stacking. Pseudouridine has a higher hydrogen bonding capacity than uridine and increases the rigidity of the sugar-phosphate backbone (84, 85). m^6^A-modified RNA is recognized by specific proteins (86). These chemical and topological properties may affect critical RNA-RNA or RNA-protein interactions. Because of the sequence diversity of the SL in *A. carterae*, further investigations into the specific base modifications and arrangement within every possible SL have not been pursued. However, given sequence evidence of heterogeneity within the SL and a range of size products after incubation with T2 nuclease, it seems likely that base modifications within the SL may also be heterogeneous and may play a role in RNA recognition by eIF4E family members.

### eIF4E-1a emerges as the likely translation initiation factor for dinoflagellates

eIF4E is part of an extended gene family found exclusively in eukaryotes. Throughout the eukaryotic domain, a series of eIF4E gene duplications has led to the generation of a family of proteins with multiple structural classes and in some cases subclasses within a given organism. The translation factor, eIF4E, is defined by the “cupped-hand” structure within which the mRNA cap is bound, as exemplified by the mouse eIF4E1A, a prototypical metazoan Class I cap binding translation initiation factor. The mRNA cap-binding region is found within a core of 160 to 170 amino acids containing eight aromatic residues with conserved spacing (40). The secondary structure consists of six beta sheets which line the binding pocket and three major alpha helices (84, 85, 87, 88). In mouse eIF4E1A, recognition of the 7-methylguanosine cap base is mediated by base sandwich-stacking between conserved residues W56 and W102 as well as with E103. In addition, W166 interacts with the methyl group on the modified base of the mRNA cap. Furthermore, the triphosphate of the cap base forms salt bridges with R112, R157 and K162 (87, 88). The alpha helices form the exterior, solvent accessible side of the protein with alpha helix one, containing a recognition motif of S/TVxxFW that interacts with eukaryotic translation initiation factor 4G (eIF4G) and a range of unrelated eIF4E-interacting proteins (38, 89).

Phylogenetic analysis of eIF4E family members in core dinoflagellates revealed between eight and fifteen members per species composed of three clades, eIF4E-1, eIF4E-2 and eIF4E-3 and nine distinct subclades (32) which cannot be mapped to the previously described metazoan eIF4E classes, Class I, Class II and Class III (32, 40). The large sequence divergence between the three dinoflagellate clades implies three different roles. This is supported by the finding that the divergencies between the three clades are found at amino acids critical for initiation factor function suggesting the family members are functionally distinct. Initial sequence comparisons of clade 3 eIF4E family members from *A. carterae* and *Karlodinium veneficum* indicated many attributes in common with metazoan Class 1 eIF4Es; a higher identity/similarity to metazoan Class I eIF4Es than eIF4E family members from clades 1 and 2, a tryptophan at the position equivalent to W56 in mouse eIF4E1A and complete conservation of the amino acids important in charge neutralization of the phosphate bridge of the mRNA cap. However, clade 3 eIF4Es are not found in ciliates or apicomplexans. Furthermore, they do not bind cap, are barely detectable and do not support growth in yeast conditionally dependent on eIF4E. This observation was a good lesson in not relying on homology to assume function.

Overall, eIF4E-1a emerges with characteristics consistent with the role of a prototypical initiation factor; eIF4E-1a is the most conserved and highly expressed eIF4E family member, has the highest affinity for m^7^GpppG and m^7^GpppC by surface plasmon resonance, and is able to complement a yeast strain conditionally deficient in eIF4E. Furthermore, eIF4E-1a has potential phosphorylation sites equivalent to Ser209 in mouse/human eIF4E1A. Although phosphorylation of Ser209 in mouse/human eIF4E1A reduces overall eIF4E–mRNA association it results in preferential association or dissociation of subsets of mRNAs (90).

The translation initiation factor, eIF4E, recruits mRNA to the ribosome through its ability to bind the 5’ cap structure. However, eIF4E has no means to bind the ribosome itself but, in metazoans, does so through its interaction with the large scaffold protein, eIF4G, an eIF4E binding protein (reviewed (38, 91). eIF4G possesses a modular structure with binding sites for different initiation factors that is highly conserved in metazoans. in particular, its central region, MIF4G/HEAT1 domain, an eIF4E-binding site and a poly(A) binding protein (PABP) binding site. In eIF4E family members capable of binding eIF4G and other eIF4E interacting proteins, the consensus sequence of the recognition motif is S/TVxxFW ending at Trp-73 in mouse eIF4E1A. In *A. carterae* eIF4E family members, similar motifs are conserved. Although clade 1 and −3 eIF4E family members from *A. carterae* have a tryptophan at the position equivalent to Trp-73, eIF4E-2 has phenylalanine. In addition, there are subtle variations between the different eIF4E-1 sub-clades in the consensus sequence of the binding domain. The region can be polar (TVQEFW), as in eIF4E-1a and -1b, acidic (TVEEFW) as in eIF4E-1c, or basic (TVKGFW), as in eIF4E-1d suggesting interaction with different binding partners that could result in interaction with different subsets of mRNAs.

The role of these motifs in dinoflagellates is unclear particularly since prototypical eIF4G has not been identified in dinoflagellates. Although there are several candidate genes that have eIF4G conserved domains, they are significantly different from canonical metazoan eIF4Gs in that they lack the amino-terminal eIF4E and PABP binding domains (37). All dinoflagellate eIF4G transcripts found to date encode only that portion of metazoan eIF4G equivalent to the MIF4G/HEAT1 domain. This form is expected to allow interactions with the translation factors eIF4A, eIF3 as well as RNA. In metazoan eIF4Gs, the binding domains for eIF4E and PABP allow mammalian eIF4G to coordinate independent interactions with mRNA via the cap and poly(A) tail which promotes circularization of the mRNA and a more efficient re-utilization of recruited mRNAs (92). The shortness of the *A. carterae* eIF4G candidates may only reflect the difficulty of identifying full length transcripts that encode such large proteins. Alternatively, other translation components may contribute to the interaction of eIF4E and eIF4G. However, a variety of alternative mechanisms to initiate translation in the absence of eIF4G have been uncovered (reviewed (93). Alternate eIF4E binding proteins, unrelated to eIF4G are able to serve as the central scaffold. These include threonyl tRNA synthetase in vertebrates (94), mextli in *Drosophila melanogaster* (95), CERES in *Arabidopsis thaliana* (96) and eIF3a in *Leishmania major* (97).

Like dinoflagellates, trypanosomatids have atypical shorter eIF4Gs that lack eIF4E or PABP binding domains. Nevertheless, interaction between these eIF4Gs and the trypanosomatid EIF4Es to form activated mRNP complexes has been shown (reviewed in (41, 47). Trypanosomatid genomes encode six divergent orthologues of eIF4E (EIF4E1 to EIF4E6) that fall into three different clades, with a low degree of conservation among themselves (sequence identity below 22 %, except for EIF4E5 and EIF4E6, with ∼30 % of sequence identity), and that are significantly different from their counterparts from plants and metazoans. Trypanosomatid EIF4E3 and EIF4E4 are considered to be the canonical translation factors required for cap4-dependent translation (41, 47). Five orthologues of eIF4G are found in Trypanosoma and Leishmania species. While *Leishmania* EIF4E3 interacts with eIF4G4, eIF4G3 interacts with EIF4E4. The multiple truncated, atypical eIF4Gs of trypanosomatids have been shown to associate with different eIF4Es, as well as eIF4As and PABPs to support translation. For instance, *Leishmania* eIF4E4 is capable of binding directly to PABP1 via three conserved motifs located in its amino-terminal domain potentially, allowing an unusual cap4-eIF4E-PABP1-poly(A) bridge to circularize mRNAs involved in translation (41).

Throughout eukaryotes, not only are there multiple eIF4E family members with a range of functions, but a range of eIF4E-interacting proteins that direct their activity in translational initiation or translational repression, in pre-mRNA processing, nucleocytoplasmic mRNA export or mRNA storage in cytoplasmic foci (reviewed Hernández, 2022; Hernández and Vazquez-Pianzola, 2023, #46247}. These include proteins such as the 4E-BPs, maskin, cup, 4E-T, neuroguidin, LRPPR1, and others (reviewed (38, 89)). The only putative eIF4E-binding protein identified so far in dinoflagellates is neuroguidin, which has three eIF4E binding motifs, and is found also in both *Lingulodinium* and *Symbiodinium* spp (e-values of e^-15^). In the axons and dendrites of the mammalian nervous system, neuroguidin binds to eIF4E1A and represses translation of cytoplasmic polyadenylation element (CPE)-containing mRNAs (98), but it is not known what role, if any, they play in dinoflagellates. No sequences encoding the regulatory eIF4E binding proteins (4E-BPs) have been identified in dinoflagellates to date. However, 4E-BPs are not consistently found throughout the eukaryotic tree of life. They are absent from trypanosomatids, plants and nematodes, but are present the fungi Basidiomycetes, and the unicellular taxa Amebozoa, Glaucocystophyta, and Jakobids (99).

### Could the large number of eIF4E family members in *A. carterae*, along with the diverse and heavily modified spliced leader sequences account for the regulation of mRNA recruitment seen in dinoflagellates?

There is no simple and uniform system describing the function and structure of eIF4Es in all eukaryotes and particularly in protists. Trypanosomatids have six eIF4E family members, higher than that of metazoans (41) and dinoflagellates have even more (eight to fifteen). For trypanosomatids, this was attributed to their complex life cycle as parasites with multiple hosts and several differentiation stages and has been thought to allow for reprogramming of protein expression when facing varying conditions in different hosts (41, 100). Dinoflagellates and trypanosomatids share a lack of transcriptional regulation and also share SL *trans*-splicing. Both lineages share m^7^G as the 5′-cap base. Trypanosomatids have modifications in the first four cap proximal bases, referred to as cap-4. In contrast, *A. carterae* has a large number of eIF4E family members along with diverse SL sequences and heavy modification of its mRNA SLs. These features underscore the unique nature of the translational machinery in the dinoflagellate lineage and point to a wide range of possibilities for differential recruitment of mRNAs to the translation machinery.

The extent of the regulation of mRNA recruitment in dinoflagellates has been made clear recently in *Lingulodinium polyedrum*. Like other dinoflagellates, *L. polyedrum* show many metabolic changes over a diel period, although few differences in transcript abundance can be detected (37, 101). Using a simplified method of ribosome profiling, Morse et al have been able to document genome-wide changes in RNA translation rates over the diel cycle (102–104). Their studies have identified over three thousand transcripts that show regulated recruitment to ribosomes over the course of a daily light-dark cycle. However, recruitment is nuanced and complicated such that transcripts involved in the same metabolic processes are coordinately recruited.

In *L. polyedrum*, there are three different times at which large numbers of transcripts have peak translation rates, with roughly a third having their maximum recruitment rates at the dark/light interface (termed ZT0), a third at the light/dark transition (ZT12) and a third around ZT16. Transcripts expressed at a single time are clustered into particular biochemical pathways and represent very different groups of transcripts. All the transcripts of enzymes involved in photosynthesis are concurrently recruited at the light/dark interface, ZTO, predicting the known rhythm of carbon fixation. Transcripts encoding enzymes in the glycolytic pathway also show peak recruitment at ZT0. At ZT12, transcripts encoding components of bioluminescent reactions show peak recruitment as do transcripts of many enzymes involved in DNA synthesis and purine and pyrimidine metabolism. At ZT16, a third peak in recruitment is seen, encompassing transcripts encoding translation factors, ribosomal proteins and enzymes involved in amino acid biosynthesis.

This unique, highly coordinated, recruitment of transcripts in dinoflagellates could be explained by the unique nature of the translational machinery; the large number of eIF4E family members, the possibility of regulation by phosphorylation of eIF4E-1a, along with diverse spliced leader RNA sequences and heavy modification of the mRNA spliced leaders. These features point to a wide range of possibilities for differential recruitment of mRNAs to the translation machinery.

## Materials and Methods

### Cell culture and harvest

*Amphidinium carterae* Hulburt strain CCMP 1314 was isolated in 1954 by Robert Guillard from the Falmouth Great Pond, Falmouth, Massachusetts USA (41.56° N 70.583° W). The strain was obtained from Bigelow National Culture of Marine Algae and Microbiota as an axenic culture (https://ncma.bigelow.org/ ccmp1314#.Vu8OHBIrIo8). Cells were grown in ESAW-32 medium (105) modified to contain 10 mM HEPES, pH 8.0 and 32 ppt saline content, in a 20 L multiport polycarbonate carboy. *Karlodinium veneficum* NCMA strain number 2936 (https://ncma.bigelow.org/ccmp2936#1.Vu8ORxIrIo8) was cultured in the same medium that was diluted with fresh sterile water to 15 ppt. The cultures were maintained under a light intensity of 150 μm cm^-2^ s^-1^ with a 14:10 light:dark schedule and bubbling of air infused with CO_2_ controlled by an American Marine (Ridgefield Ct) pH controller set to turn on above pH 8.2 and off at pH 7.6. Two liters of medium were added to an actively growing culture on Monday, Wednesday, and Friday until reaching a cell density of 172,000 cells/ml on the morning of the experiment. 250 ml of culture was aliquoted into 26-75 cm2 polystyrene culture flasks from Corning (Corning NY) and placed along the light bank to achieve equivalent light exposure to the stock culture using two rows of thirteen, with duplicates in opposing directions. Each duplicate pair was taken for harvest according to the following schedule relative to the transition from light to dark in hours: -6.0 and +6.0. The samples were each split into 50-ml and 200-ml aliquots. Each aliquot was centrifuged at 1000 x g for 10 min at 20° C. The 200-ml aliquot was frozen at -80° C and the 50-ml aliquot was suspended in 1 ml of TRI reagent (Sigma-Aldrich Saint Louis, MO) for RNA isolation. Cell abundance and equivalent spherical diameter (ESD) were measured on a Coulter® Counter (Beckman Coulter, Fullerton, CA USA) using the ‘narrow’ size range (4–30 μm). Equivalent spherical volume (ESV) was calculated as (4/3)(πr3), where r is the radius derived from ESD. A 100 μl sample was counted. Samples that yielded a ‘coincidence correction’ value above 20 % were diluted 1:10 with sterile seawater at a salinity of 32 ppm.

### RNA extraction and RT-qPCR analysis

Duplicate RNA samples from each time point were extracted using TRI reagent according to the manufacturer’s protocol using 1 ml of TRI reagent. RNA was quantified on a Nanodrop 1000 (Thermo Fisher) and also on a Qubit 2.0 fluorometer (Life Technologies), to confirm accurate RNA abundance (data not shown). RNA was reverse transcribed using Superscript II Reverse Transcriptase (Life Technologies) with random primers (Invitrogen) according to the manufacturer’s protocol. Generated cDNA was used as template for quantitative Real-Time PCR using an Applied Biosystems (Life Technologies) Fast 7500 thermal cycler. Primers for each eIF4E family member were designed using Primer3 (106). Two primer pairs were designed for some to target different areas of the mRNA to confirm the results (Fig. 6<u>-Suppl. Table 1</u>). Primers were designed to function effectively in a qPCR measurement experiment yielding a DNA product of approximately 150 bp.

Each reaction was performed in duplicate with the following setup: 6 μl diethyl pyrocarbonate (DEPC) treated water, 2 μl of combined forward and reverse primers at 5 μM each, 10 μl of iTAQ 2X master mix containing SYBR green and ROX (Bio-Rad Hercules, CA), and 2 μl of template cDNA at 10 ng/μl. Thermal cycling conditions consisted of an initial denaturation at 95 °C for 2 min followed by 40 cycles of denaturation at 95° C for 15 sec, annealing and fluorescent data collection at 60° C for 15 sec, and extension at 72° C for 30 sec. The reaction was completed with a melt curve to determine the presence of spurious PCR products. Cycle thresholds and baselines were determined manually and relative quantities were determined across the two diel time-points and expressed relative to the large ribosomal targeted product and assumed primer efficiency of 1.85 copies/cycle. This was done for normalization across the time points and not to produce accurate quantities, thus primer efficiencies were not determined empirically for each primer pair. Expression levels were normalized to total input RNA from each time point.

### Enrichment of Spliced Leader-Containing Sequence and 5′ Cap Isolation

Twelve 4-liter samples of an exponentially growing *A. carterae* culture were taken twice weekly, pelleted at 1000 x g for 10 min and frozen at -80° C. Each pellet was used for RNA extraction using the RNAzol RT reagent (Molecular Research Center, Cincinnati, OH) according to the manufacturer’s instructions. Briefly, pellets were homogenized in the RNA extraction reagent, the nucleic acids were allowed to deproteinate at room temperature for 15 min, and cellular debris was removed by centrifugation. Large RNAs (approximately >200 bases) were precipitated with 0.4 volumes of 75 % ethanol and pelleted by centrifugation. The remaining small RNA fraction was precipitated from the supernatant with 0.8 volumes of isopropanol and pelleted by centrifugation. The pellets were suspended in water treated with diethyl pyrocarbonate (DEPC) and quantified. The isolated RNA from each size fraction was pooled and re-precipitated with an equal volume of isopropanol at room temperature for 15 min, pelleted at 12,000 x g for 20 min, washed with 75 % isopropanol, and re-pelleted at 12,000 x g for 5 min. The final pellets were suspended in 200 μl of DEPC water, yielding approximately 200 μg of RNA >200 bases and 5.2 μg of RNA <200 bases.

For the U4 and SL pulldowns, 1 μg of the small RNA fraction was diluted to 44 μl and combined with 5 μl of 5 M NaCl and 1 μl 1 M MgCl_2_, each. The RNAs from the >200 base fraction were similarly diluted and also used for spliced leader enrichment. 50 μl of formamide was added to each 50-μl sample and combined with 100 μl 2x hybridization buffer (8X SSC, 1mM EDTA, 20 % dextran sulfate). 400 μl of streptavidin coated beads were washed with washing buffer and bound to the biotinylated primers for 1 h at room temperature on a rotisserie. The RNA samples in hybridization buffer were combined with 100 μl of the bead bound oligonucleotide and hybridized for 18 h at 40° C. The beads were washed five times with 500 μl washing buffer. The SL pulldown from the large RNA fraction was suspended in 100 μl RNAse A buffer (10 mM Tris-HCl, pH 7.6, 1 M NaCl). The SL pulldown from the small RNA fraction was suspended in 100 μl of RNAse T2 buffer (10 mM ammonium acetate, pH 4.5, 10 mM EDTA), and the U4 sample was suspended in 100 μl decapping buffer. 1 μl of 10 mg/ml RNAse A (Thermo, Waltham, MA) was added to the large RNA SL pulldown and single stranded RNA was degraded at 37 °C for 1 h. 1 μl recombinant RNAse T2 (Mo Bi Tech, Goettingen, Germany) was added to the small RNA SL pulldown and incubated at 37° C for two h to achieve complete digestion. Following RNAse T2 treatment, the SL pulldowns were each washed five times with 500 μl washing buffer and suspended in 100 μl decapping buffer. The RNA from the U4 and the two SL pulldowns was melted off the bead-bound oligonucleotide at 70° C for 5 min and separated immediately from the beads. 1 μl of each sample was then imaged on the Agilent Bioanalyzer 2100 using the small RNA kit to verify product sizes. 2.5 μl of DCP2 (Enzymax, Lexington, KY) was added to each SL pulldown and decapping was performed at 37° C for 30 min. The 22-base presumed spliced leader from the large RNA fraction was removed from the excised 5′ cap by addition of 100 μl of decapping buffer containing streptavidin coated beads bound to the spliced leader complementary oligo and annealing of the presumed spliced leader to the oligo at 45° C for 5 min. The beads were removed, suspended in 100 μl of decapping buffer, and the presumed spliced leader was melted off the oligo at 70° C for 5 min and the beads removed. The SL pulldown from the small RNA fraction was transferred to a 3000 NMWL regenerated cellulose Amicon spin filter (Millipore, Billerica, MA). The removed 5′ cap was enriched by increasing the sample volume with 500 μl decapping buffer and passing the sample through the filter at 10,000 x g twice. The resultant samples from the U4 pulldown and the isolated caps and decapped substrates from the two SL pulldowns were lyophilized and prepared for compositional analysis.

### Compositional analysis of purified RNAs

Each purified RNA sample was aliquoted at a final concentration of 0.73 ng/µl, 7.3 ng in 10 µl of RNAse free water and 10 pg/µl, 100 pg in 10 µl, of isotopically labelled [^13^C][^15^N]-guanosine as internal standard. Each individual sample underwent two-step enzymatic hydrolysis followed by ultra-high performance liquid chromatography (UHPLC) tandem mass spectrometry (MS/MS) method as previously described. The first part of the digestion involves an endonucleolytic cleavage to yield 5’-phosphate with nuclease P1. One unit of nuclease P1 from *Penicillium citrinum* (Sigma-Aldrich) was added to each sample and incubated overnight at 37 °C. The second step of the hydrolysis was performed by the addition of a unit of bacterial alkaline phosphatase from *E. coli* (Sigma-Aldrich) at 37 °C for 2 h. This enzyme specifically cleaves the 5’-phosphate from the nucleoside resulting in individual nucleosides and inorganic phosphate. The nucleoside products were lyophilized and reconstituted in 40 μl of RNAse free water (18.0 MΩcm-1) containing 0.01 % formic acid prior to UHPLC-MS/MS analysis.

Nucleoside mixture products from hydrolysis were subject to chromatographic separation on a Waters Acquity I-Class UPLCTM (Waters, USA) equipped with a binary pump and auto-sampler maintained at 4 °C. A Waters Acquity UPLCTM HSS T3 guard column (2.1 x 5 mm 1.8 µm) followed a HSS T3 column (2.1 x 50 mm 1.7 µm). Column temperature was set at 25 °C. The mobile phases included RNAse-free water (18.0 MΩcm-1) containing 0.01% formic acid pH 3.5 (Buffer A) and 50 % acetonitrile in aqueous 0.01 % formic acid (Buffer B). Flow rate was set up at 0.2 ml/min and a gradient applied as described previously.

Tandem MS analysis of nucleosides provides a second-dimension analysis whereby the induction of collision energy as the protonated molecular ion [MH+] passes through the collision cell will produce a specific secondary ion or product ion [BH2+]. Generally, the protonated nucleoside, molecular ion [MH+], is fragmented at the glycosidic bond providing the protonated nucleobase, product ion [BH2+] and the neutral sugar residue. Tandem MS analysis was performed on a Waters XEVO TQ-STM (Waters, USA) triple quadrupole mass spectrometer equipped with an electrospray ionization (ESI) source maintained at 150 °C and the capillary voltage was set at 1 kV. Nitrogen was used as the nebulizer gas which was maintained at 7 bars of pressure, flow rate of 500 l/h and a temperature of 500 °C. UPLC-MS/MS analysis was determined in ESI positive-ion and multiple reaction monitoring (MRM) mode with retention times and corresponding molecular and product ion pairs [MH+]/[BH2+] as input parameters.

Quantitation of the 35 nucleosides, 4 majors and 31 modifications, was performed using standard curves with concentration ranging between 0.05 and 100 pg/ul of pure nucleosides and isotope-labelled internal standard [^13^C][^15^N]-guanosine. Concentrations of nucleosides were calculated following Beer-Lambert law where the extinction coefficient was itself calculated at the absorbance maximum nearest 260 nm rather than as the standard 260 nm as most of the nucleosides have peak maxima that differs from that at 260nm (107).

To determine the presence of RNA modifications from RNA extracts and to quantify them, we performed UHPLC-MS/MS measurements. A negative control sample was included in the data set to account for background signal from possible modifications part of enzyme’s background that could interfere with nucleosides of interest. The negative control contained the enzymes, internal standard and reagents used during the enzymatic digestion. After digestion each sample was lyophilized and reconstituted in RNAse free water (18.0 MΩcm-1) containing 0.01 % formic acid to a final concentration of 180 pg/µl of RNA and 1 pg/µl internal standard. A blank sample containing water in 0.01 % in formic acid solution was analyzed between each sample to avoid cross-contamination. Each sample type included three biological replicates and three technical replicates to account for instrument variability. From the thirty-one RNA modifications included in the UHPLC-MS/MS method, ten were above the limit of detection, including mostly methylations and pseudouridine. Those modifications below the limit of detection were excluded from the results. Since it was not possible to take total RNA through the streptavidin-based enrichment process to use as a negative control, modified RNAs detected in the sample that contained the isolated m^7^G cap were used to normalize other samples. Since these other residues are theoretically degraded RNA moieties carried through the enrichment process, they were our best estimate for the levels of modifications in the total RNA pool used, mainly ribosomal and mRNAs.

### Identification of *A. carterae cap* structure and RNA modifying genes

BLAST searches to identify contigs with the necessary domains to perform the capping reactions were performed as an early screen in the *A. carterae* transcriptome. Candidate contigs were used as queries to retrieve similar sequences from other transcriptomes, and a phylogenetic tree was created using these and apicomplexan sequences to verify the reconstruction of the organismal phylogeny.

### Identification of *A. carterae* eIF4E family members

The *A. carterae* eIF4E sequences were from a previous analysis of eleven dinoflagellate transcriptomes (32). eIF4E sequences were retrieved and annotated by initial query with the mouse eIF4E-1 amino acid sequence, NP_031943.3. The resulting eIF4E sequences were aligned by the MUSCLE algorithm, using MacVector (MacVector Inc.). The alignments in this manuscript were colored with the BOXSHADE web application (http://www.ch.embnet.org/software/BOX_form.html) and stylistically edited in Adobe Illustrator (Adobe). CLC Bio (Qiagen) was used to calculate the properties of each amino acid sequence. An online isoelectric point calculator (http://isoelectric.ovh.org) was used to calculate the average pI of each amino acid sequence using a variety of computational methods.

### Structure prediction and visualization

The predicted amino acid sequence of each eIF4E from *A. carterae* was analyzed using Phyre2, the Protein Homology/analogY Recognition Engine 2 (108). Using the top ten highest scoring alignments, Phyre2 constructs a 3D model of the protein structure with missing or inserted sequence modeled using the “loop library” and a reconstruction procedure. This program creates a PDB file that can be used with visualization software to view the predicted protein structure. Each PDB file was visualized using the Visual Molecular Dynamics (VMD) software package (109). The murine eIF4E model [PDB file: 1L8B], was downloaded and used to create a STAMP, structural alignment of multiple proteins, in order to position the m^7^GTP cap molecule in an approximate position to the modeled protein structure. Images were rendered using Tachyon, included with the VMD software.

### Cloning of eIF4E constructs into *in vitro*, bacterial, and yeast expression vectors

Nucleotide sequences for *A. carterae* eIF4E family members were codon-optimized for rabbit, *Oryctolagus cuniculus*, using Advanced Optimum Gene™ software (Genscript). The nucleotide sequence was synthesized by Genscript and cloned into the *in vitro* transcription/translation plasmid vector pCITE-4a(+) (Novagen), using the NcoI and BamHI sites which adds an S·tag to the amino-terminus and includes a stop codon at the carboxy terminus. Each eIF4E sequence was also sub-cloned into a re-engineered bacterial expression vector pGEX-4T (GE Healthcare), now pGEX-4T(MCS)-TEV, in which the multiple cloning site had been changed to include NcoI and BamHI cloning sites and a TEV protease site added between the GST-tag and the cloned gene. In addition, each eIF4E sequence was codon-optimized for *S. cerevisiae*, synthesized by Genscript, and cloned into the yeast expression vector, pRS416GPD (ATCC), using the BamHI and XhoI sites, respectively.

### Bacterial expression of recombinant eIF4E family members

Each pGEX-4T(MCS) construct containing an eIF4E gene was transformed into JM109 competent *E. coli* (Promega) and grown on LB + 100 μg/ml carbenicillin (Sigma). Purified plasmid constructs were used to transform the bacterial expression strain Rosetta™(DE3)pLysS (EMD Millipore). To express protein, transformed cells were grown in 500 ml of autoinduction media (110), supplemented with 34 μg/ml chloramphenicol and 100 μg/ml carbenicillin, at 25 °C over three days with shaking at 300 RPM. On the third day, cultures were pelleted at 20,000 x g and washed with phosphate buffered potassium, pH 7.5, before being frozen at -80 °C. Frozen pellets were suspended in lysis buffer (20 mM phosphate, pH 7.5, 150 mM KCl, 1x Protease Inhibitor Cocktail (Pierce). After thorough suspension, the bacteria were lysed using a chilled French Press, pressurized to 20,000 PSI. The resulting lysate was clarified by ultracentrifugation at 25,000 x g for 20 min at 4°C.

### Generation and validation of affinity-purified antibodies

The amino acid sequences of each eIF4E family member, except eIF4E-1d2 and -3b, from *A. carterae* were submitted to the Genscript Optimum Antigen™ Design Tool to determine the optimal antigenic regions to use for immunization. Genscript synthesized each antigenic peptide (Fig. 7<u>-Suppl. Table 3</u>) and added an additional cysteine residue to allow for conjugation to the KLH adjuvant. These were used for immunization of New Zealand white rabbits. Specific antibodies were isolated from the resulting serum by affinity purification using the synthesized peptide as bait. Antibodies were tested for reactivity by an ELISA assay with the peptide used to generate the antibody as the target coating the wells of a 96-well plate.

Antibodies were also tested for specificity and reactivity using western blot with recombinant protein. In brief, each GST-tagged recombinant protein was purified either by GST column chromatography or by inclusion-body purification using protein solubilized after denaturation with urea and buffer exchange. Protein concentrations were determined using a Qubit™ protein assay kit (Life Technologies) and approximately 250 ng of each protein loaded per lane. Samples were analyzed using a 4-12 % Bis-Tris NuPAGE SDS-PAGE gel (Life Technologies) and separated by molecular weight using denaturing electrophoresis. Proteins were transferred to PVDF 0.2 μM membrane (BioRad) using the Bolt Mini Blot Module (Life Technologies) and western blot carried out using the iBind Western Blot Apparatus (Life Technologies). Affinity-purified rabbit polyclonal antibodies specific to each eIF4E family member were used as the primary probe in western blotting. An anti-rabbit IgG (H&L) (GOAT) antibody that was peroxidase conjugated (BioRad) was used as the secondary probe. Luciferase signal was visualized by incubation in Clarity Enhanced Chemiluminescent (ECL) substrate (BioRad) and imaged using a FluorChem E (Protein Simple). Quantification of band intensities was performed using Imageview software (Protein Simple).

### Quantification of eIF4E protein levels from *A. carterae*

Protein levels for each eIF4E family member were assessed by western blot analysis using whole cell lysates solubilized into SDS-sample buffer. Western blot analysis was used to quantify the relative levels of each eIF4E family member from *A. carterae* for which an antibody and purified protein (>90 % pure) was available. Each recombinant protein concentration was quantified and diluted into a two-fold standard curve either starting with 25 ng or 3 ng total protein loaded per lane of an SDS-PAGE gel. An actively growing mid-day culture of *A. carterae* was quantified for cellular concentration and centrifuged to yield a pellet of known cell number. This was suspended in 1X SDS-Sample buffer and boiled so that when diluted into a two-fold series it would start with either 2×10^5^ or 5×10^5^ cells/lane. Some eIF4E family members were present at such a low abundance compared to the others that the standard curve or cell equivalents had to be adjusted simultaneously to allow for the eIF4E in question to be visualized in the western blots. The dilutions for both cell number per lane and protein concentrations were taken in to account when molecules per cell were calculated.

### Mass spectrometry reveals three of the eIF4E family members isolated from a 27-33 kD protein fraction from *A. carterae* whole cell lysates

Gel bands corresponding to the 27-33 kDa location, based on the SeeBlue2 plus molecular weight ladder (Thermo Fisher Scientific), were excised with a clean scalpel. Gel bands were destained twice with 200 μl destaining solution (∼25 mM sodium bicarbonate in 50 % acetonitrile) and incubated at 37 °C with shaking for 30 min. Samples were processed using the In-Gel Tryptic Digestion Kit (Thermo Scientific, Inc.) according to manufacturer’s protocol. The peptides generated were analyzed by electrospray ionization on an Elite tandem orbitrap mass spectrometer (Thermo Scientific Inc). Nanoflow HPLC was performed by using a Waters NanoAcquity HPLC system (Waters Corporation). Peptides were trapped on a fused-silica pre-column (100 μm i.d. 365 μm o.d.) packed with 2 cm of 5 μm (200 Å) Magic C18 reverse-phase particles (Michrom Bioresources, Inc.). Subsequent peptide separation was conducted on a 75 μm i.d. x 180 mm long analytical column constructed in-house and packed with 5 μm (100 Å) Magic C18 particles, using a Sutter Instruments P-2000 CO2 laser puller (Sutter Instrument Company). The mobile phase A was 0.1 % formic acid in water and mobile phase B was 0.1 % formic acid in acetonitrile.

Peptide separation was performed at 250 nl/min in a 95-min run, in which mobile phase B started at 5 %, increased to 35 % at 60 min, 80 % at 65 min, followed by a 5-min wash at 80 % and a 25-min re-equilibration at 5 %. Ion source conditions were optimized by using the tuning and calibration solution recommended by the instrument provider. Data were acquired by using Xcalibur (version 2.8, Thermo Scientific Inc.). MS data were collected by top-15 data-dependent acquisition. A full MS scan of range 350–2000 m/z was performed with 60 K resolution in the orbitrap followed by collision induced dissociation (CID) fragmentation of precursors in the iontrap at normalized collision energy of 35 %. The MS/MS spectra of product ions were collected in rapid scan mode.

Acquired tandem mass spectra were searched for sequence matches against the UniprotKB database and custom 6-frame translated transcriptome database for *A. carterae* using COMET. The following modifications were set as search parameters: peptide mass tolerance at 10 ppm, trypsin digestion cleavage after K or R (except when followed by P), one allowed missed cleavage site, carboxymethylated cysteines (static modification), and oxidized methionines (variable modification/differential search option). PeptideProphet and ProteinProphet, which compute a probability likelihood of each identification being correct, were used for statistical analysis of search results. PeptideProphet probability 0.9 and ProteinProphet probability 0.95 were used for positive identification at an error rate of less than 1 %.

### *In vitro* transcription/translation of eIF4E and luciferase constructs

*In vitro* translation (IVT) of each eIF4E was performed using the TNT Quick Coupled Transcription/Translation System (Promega) according to the manufacturer’s protocol and based on a protocol developed to study zebrafish eIF4E (111). In brief, approximately 1 μg of plasmid DNA and 20 μCi of Easy-tag ^35^S-methionine (Perkin Elmer) were mixed with a final volume of 50 μl, in which TNT constituted 80 %. Transcription and translation were carried out at 30 °C for 1.5 h. 1 μl of the IVT reaction was taken for analysis of ^35^S-methionine incorporation by mixing to a final concentration of 5 % TCA, boiling and capturing on GF/C filter paper. The filter paper was used to analyze CPM incorporation in a scintillation counter.

### m^7^GTP- and trimethyl-guanosine triphosphate-Sepharose binding assay

Sepharose beads bound to 7-methyl-guanosine-triphosphate (Jena) and trimethyl-guanosine triphosphate (kindly provided by Edward Darzynkiewicz) were blocked using 1 mg/ml soybean trypsin inhibitor (Sigma, T9128) in binding buffer (25 mM HEPES/KOH pH 7.2, 10 % glycerol, 150 mM KCl, 1 mM dithiothreitol, 1 mM D-L methionine) for 1 h at 4° C with shaking at 1400 rpm in a benchtop thermomixer 22331(Eppendorf). The beads were washed twice with binding buffer without soybean trypsin inhibitor and suspended in 50 % v/v binding buffer. 20 μl of each IVT reaction was diluted 10-fold with binding buffer containing 200 μM GTP and 200 μM MgCl^2^ and mixed with the bead suspension and incubated at 4° C for 1 h with shaking at 1400 rpm. The supernatant containing the unbound fraction was recovered by centrifugation at 500 x g at 4° C. An equivalent of 1 μl of the original IVT was used for TCA precipitation and filtered onto a GF/C membrane (Millipore). The cap-analogue beads were washed 5 times with binding buffer and the final bead fraction suspended in SDS-PAGE sample buffer. The bead suspensions were heated to 90° C and a fraction equivalent to 1 μl of the original IVT reaction was applied to GF/C filter paper. Fractions were counted in Ecoscint Original scintillation cocktail (National Diagnostics, Georgia, USA) and ^35^S was determined using a LS6500 Multipurpose Scintillation Counter (Beckman Coulter). IVT, unbound, and bead bound fractions were diluted in SDS-PAGE sample buffer and heated to 90° C for 3 min. The samples were separated by 17.5 % high-Tris SDS-PAGE using PAGEruler pre-stained molecular weight ladder (Fermentas) as a guide and transferred to Immun-blot PVDF (BioRad) using a Criterion blotter (BioRad) for 30 min at 100 V in 20 % methanol Towbin buffer (BioRad). Labeled proteins were visualized on the PVDF membrane using a Storage Phosphor screen (Molecular Dynamics) and imaged with a Typhoon 9410 Variable Mode Imager (GE Healthcare). Due to background binding and loss of protein in the washes, both fractions do not always add up to 100 %. However, this assay serves well as a qualitative approximation of protein cap binding capacity.

### Recovery of eIF4E from *A. carterae* cell extracts by m^7^GTP-Sepharose binding

Approximately 1×10^6^ cells from a mid-day growing culture of A. carterae were harvested by centrifugation at 1,000 × g for 10 min. The cell pellet was flash-frozen with an ethanol/dry ice bath. The pellet was suspended in 2 ml of lysis/binding buffer (25 mM HEPES/KOH, pH 7.6, 10 % glycerol, 150 mM KCl, 1 mM dithiothreitol, 0.1 % Elugent (Calbiochem), loaded into SHREDDER tubes (Pressure Biosciences Inc; Easton, MA) and disrupted using a Barocycler NEP2320 at 35,000 PSI for 1 min for 30 cycles. The lysate was clarified by centrifugation at 10,000 x g for 10 min at 4° C and the insoluble fraction suspended in 1x SDS-PAGE sample buffer. The supernatant was removed and incubated with m^7^GTP-Sepharose beads (Jena Biosciences) with shaking at 1,200 RPM and 4° C for 1 h using a temperature-controlled Thermomixer 22331 (Eppendorf). The beads were pelleted at 500 x g for 1 min and the supernatant removed to become the unbound fraction. The beads were washed 5 times with binding buffer and bound protein eluted using 1x SDS-PAGE sample buffer and boiled to prepare for electrophoresis. The unbound and first wash fraction were TCA precipitated and suspended in 1x SDS-PAGE sample buffer to an equivalent volume to match the bound fraction.

### Analysis by SDS-PAGE and western blotting

Equivalent volumes of each fraction were loaded onto a 4-12 % Bis-Tris NuPAGE SDS-PAGE gel (Life Technologies) and separated by molecular weight using denaturing electrophoresis. Proteins were transferred to PVDF 0.2 μM membrane (BioRad) using the Bolt Mini Blot Module (Life Technologies) and western blot carried out using the iBind Western Blot Apparatus (Life Technologies). Affinity-purified rabbit polyclonal antibodies specific to each eIF4E family member (Genscript) were used as the primary probe in western blotting. The secondary antibody was a goat anti-rabbit antibody conjugated to horseradish peroxidase (BioRad). Bound antibody was visualized by incubation in Clarity ECL substrate (BioRad) and imaged using a FluorChem E (Protein Simple). Quantification of band intensity was performed using Imageview software (Protein Simple).

### Complementation of a *S. cerevisiae* strain expressing a glucose-repressible *H. sapiens eif4E* gene

cDNA encoding the *A. carterae* eIF4E family members (-1a, -1b, -1c, -1d1, -1d2, -2a, and -3a), each cloned separately into the yeast expression vector, pRS416GPD, were transformed into *S. cerevisiae* strain JOS003 using the Quick and Easy Transformation Mix (Clontech). JOS003 is a strain in which the one endogenous EIF4E gene has been replaced by homologous recombination with a KanMX4 cassette. The transformed yeast was spot plated on synthetic deficient (SD) media lacking uracil and leucine and containing 200 μg/ml G418, with either galactose or glucose. Growth was compared to two positive controls, *D. rerio* eIF4E1A and *S. cerevisiae* eIF4E, and two negative controls, *D. rerio* eIF4E-1B and an empty vector control. We tested the ability of each *A. carterae* eIF4E family member, except eIF4E-3b, to complement this yeast strain.

### Purification of dimeric and “active” GST-tagged eIF4E from bacterial lysates

Clarified lysate was passed over a 1-ml GSTrap Fast Flow Column (GE Healthcare) equilibrated with binding/wash buffer (20 mM phosphate pH 7.5, 150 mM KCl), using an Akta Start chromatography system (GE Healthcare) at 1ml/min at 4 °C. The column was washed with 20 column volumes of binding/wash buffer (20 mM phosphate buffer, pH 7.5, 150 mM KCl), and eluted in fifteen 1-ml fractions with elution buffer (20 mM phosphate buffer, pH 7.5, 150 mM KCl, 10 mM reduced glutathione).

It was difficult to obtain high-quality and consistent data in SPR binding affinity measurements using the protein purified solely by GST chromatography. Not only does GST-tagged eIF4E form dimers, as has been described for GST itself (112), but also larger multimers that remain soluble and can be observed in gel filtration chromatography as multiple wide peaks but were not amenable to use in affinity measurements. This has been observed by other groups where dimeric (or multimeric) forms of non-cap bound eIF4E (apo-eIF4E) can create soluble aggregates (66, 113). In addition, it has been reported that non-cap bound eIF4E is unstable in HEPES-containing buffers (66). This issue was solved by treating GST-tagged protein with 50 mM dithiothreitol for 1 h at 4° C, with shaking in a benchtop thermomixer (Eppendorf) at 1,400 RPM. The resulting solution was then centrifuged to remove protein that had fallen out of solution (about half the protein in the original mixture, data not shown). The final solution was then purified to greater than 95 % purity over a Sephadex S-300HR gel filtration column (GE Healthcare) and buffer exchanged at the same time into binding buffer (20 mM phosphate buffer, pH 7.5, 150 mM KCl) that was supplemented with 0.05 % Tween-20 (Sigma) for use in SPR analysis.

Aggregation of recombinant human eIF4E has been reported to be caused by intermolecular disulfide bonds dispersed by using dithiothreitol at 50 mM (114). This issue was solved by treating GST-tagged protein with 50 mM dithiothreitol for 1 h at 4° C, with shaking in a benchtop thermomixer (Eppendorf) at 1,400 RPM. The resulting solution was then centrifuged to remove protein that had fallen out of solution (about half the protein in the original mixture, data not shown). The final solution was then purified to greater than 95 % purity over a Sephadex S-300HR gel filtration column (GE Healthcare) and buffer exchanged at the same time into binding buffer (20 mM phosphate buffer, pH 7.5, 150 mM KCl) that was supplemented with 0.05 % Tween-20 (Sigma) for use in SPR analysis. In summary, GST-tagged proteins were purified by glutathione affinity column chromatography, dithiothreitol treatment to disperse inactive (or multimeric) complexes and gel filtration to isolate GST-fusion protein dimers of the appropriate molecular weight.

### Surface plasmon resonance (SPR)

All surface plasmon resonance experiments were performed using a Biacore T-200 (GE Healthcare). A CM5 S-series Sensor chip was used to immobilize goat-polyclonal-anti-GST antibody according to manufacturer instructions. Approximately 200 μg/ml solutions of purified and DTT treated GST-tagged *A. carterae* eIF4E-1a, eIF4E-1d1, and eIF4E-2, as well as mouse eIF4E-1, were injected at 5 μl/min over lanes 2 and 4 of the chip and loaded to approximately 2,500 Response Units (RU) in each lane. Each of the 5’ cap structure analogues (m^7^GpppG, m^7^GTP, m^7^GpppC, GpppG, and GTP) were diluted into binding buffer into a two-fold dilution series starting at 250 μM, except for m^7^GTP, which had to be further diluted to a starting dilution of 62.5 μM in order avoid a saturating response. Measurements of binding took place after two start-up cycles, then a 60-sec injection at 50 μl/min of the analyte, and dissociation allowed to occur over 60 sec.

The kinetic rates were determined by fitting the experimental data to a 1:1 interaction model between the surface bound eIF4E (B) and the cap analog analyte (A) in the injection solution, A+B↔AB. The fitting procedure used to determine the rate constants from experimental data uses numerical integration methods with an iterative approximation algorithm to find the best solution to the equations presented. The concentration of the complex formed is measured in RU by SPR response and inserted into the equation, dRdt=kaCRmax-kaC+kdR. This allows for the association and dissociation equilibrium affinities to be derived. The closeness of fit is judged by the chi-square value derived by the following equation, chi square=1..(rf-rx)2n-p, where rf is the fitted value at a given point, rx is the experimental value at the same point, n is the number of data points, and p is the number of fitted parameters. The fitting algorithm is used to minimize the chi-square value, and is judged to be complete when the difference in chi-square values between successive iterations is sufficiently small. Req=KACRmaxKAC+1 was used to calculate the equilibrium association constant from the steady-state complex formation. The value of KA is obtained by fitting a plot of Req against the concentration of the analyte (C) to this equation. The KD is calculated as the inverse of the KA, KD=1KA.

Author Contributions: Conceptualization, R.J. and A.R.P.; Bioinformatic searches and phylogenetic reconstruction, G.D.J. and T.R.B.; RNA analysis by qPCR and LC/MS, E.P.W., A.R.P., and M.B.S.; protein extraction and proteomics S.H. and A.R.P.; Recombinant protein production, western blots, and antibody pulldowns, G.D.J., R.J. and A.R.P.; cap binding assays, G.D.J. and R.J.; yeast complementation, G.D.J. and R.J.; Writing—original draft preparation, G.D.J. and R.J.; Writing—review and editing, G.D.J., E.P.W., T.R.B., A.R.P., and R.J.; Project administration, R.J and A.R.P.; Funding acquisition, R.J. and A.R.P. All authors have read and agreed to the published version of the manuscript.

## Acknowledgments

We would like to thank Mike Murphy PhD from Biacore for technical assistance, Benjamin Oyler for verification of peptide assignment and Erica Dasi for pioneering the yeast complementation assays. We also thank Dr. Edward Darzynkiewicz for Sepharose beads bound to trimethyl-guanosine triphosphate and Dr. Robert Rhoads for the IFE1 construct. This paper is Contribution No.XXXX from the University of Maryland Center for Environmental Science, and No.20xxx from the Institute of Marine and Environmental Technology.

